# Frequency-dependent fitness effects are ubiquitous in rapidly adapting microbial populations

**DOI:** 10.1101/2025.08.18.670924

**Authors:** Joao A Ascensao, Keon D Abedi, Aditya N Prasad, Oskar Hallatschek

## Abstract

In simple microbial populations, the fitness effects of selected mutations are typically assumed to be constant, regardless of mutant frequency. This assumption underpins predictions about evolutionary dynamics, epistasis, and the maintenance of genetic diversity. Here, we systematically test this assumption using beneficial mutations from early generations of the *Escherichia coli* Long-Term Evolution Experiment (LTEE). Using flow cytometry-based competition assays, we find that frequency-dependent fitness effects are the norm rather than the exception, occurring in approximately 80% of strain pairs tested. Most competitions exhibit negative frequency-dependence, where fitness advantages decline as mutant frequency increases. We demonstrate that the strength of frequency-dependence is predictable from invasion fitness measurements, which explain approximately half of the biological variation in frequency-dependent slopes. We also observe violations of fitness transitivity in several strain combinations, indicating that competitive outcomes cannot always be predicted from fitness measured against a single reference strain. High-resolution measurements reveal that frequency-dependence changes substantially within the growth cycle, with the net per-cycle effect reflecting the balance of opposing dynamics at different growth phases. Our results demonstrate that even in a model system designed to minimize ecological complexity, subtle interactions between closely related genotypes create frequency-dependent selection that can fundamentally alter evolutionary trajectories.

## Introduction

The fitness effects of most mutations in ecologically unstructured, well-mixed microbial populations are generally assumed to be constant, regardless of genotype frequency^1^. This assumption, rooted in classical population genetic theory^2–4^, underlies fundamental concepts from the distribution of fitness effects^5–10^ to predictions about fixation dynamics and the evolution of genetic diversity^11–14^. In microbial evolution experiments, fitness measurements typically treat beneficial mutations as having constant effects that drive predictable selective sweeps, with evolutionary outcomes determined heavily by the rank order of fitness advantages^15–17^.

However, this framework ignores a critical ecological reality: organisms do not evolve in isolation but interact with competitors, either directly (e.g. predation) or indirectly (e.g. modifying a shared environment). Consequently and in general, selection depends on the composition of populations^18–21^. Frequency-dependent selection, where fitness effects vary with the relative abundance of genotypes, has long been recognized as a fundamental evolutionary force in natural populations which maintains genetic diversity, countering the homogenizing effects of directional selection and genetic drift^22–30^.

Microbial evolution experiments have repeatedly uncovered frequency-dependent selection, even in simple laboratory environments where ecological complexity is thought to be minimal^1,31–33^. In this context, frequency-dependent effects are often treated as exceptional, requiring special explanation. In many of these cases the environmental heterogeneity that allows for frequency-dependent selection is not imposed by the experimenter but is generated by the evolving population itself: as lineages diverge, they modify their shared environment–for instance by excreting metabolites–and thereby construct the niches that sustain their own coexistence. Stable, negatively frequency-dependent polymorphisms arose in glucose-limited chemostats founded by a single *E. coli* clone^34,35^, typically sustained by cross-feeding on secondary metabolites such as acetate–a polymorphism that recurs so reliably that it has evolved repeatedly across independent populations^36,37^. Analogous diversification has been shown to occur in spatially structured batch culture, maintained by frequency-dependent competition among spatial niche specialists^38,39^. The *Escherichia coli* Long-Term Evolution Experiment (LTEE) is no exception: negative frequency-dependent interactions and long-lived coexistence between diverged lineages have been documented in several populations^22,27,40–44^, even though the experiment was expressly designed to minimize the opportunity for such interactions to arise^1,44^.

In mixed microbial populations composed of distinct ecotypes, resource competition is expected to play a central role in the frequency-dependent selection and eco-evolutionary dynamics, since shifts in relative abundance alter the environment in ways that feed back on fitness differences^10^. The prevailing assumption, however, is that in populations composed of a single ecotype, single point mutations have negligible effects on resource usage and thus do not generate significant frequency-dependent fitness effects. Yet, previous work^21^ revealed that the invasion fitness of a *pykF* knockout–an important beneficial target of adaptation in the early LTEE–was significantly higher than prior measurements at high frequency^45,46^. While there could have been many reasons for this discrepancy, we reasoned that one possible explanation could be the presence of negative frequency-dependent fitness effects. This prompted us to investigate how common such frequency-dependence might be among early LTEE mutations.

Here, we systematically investigate frequency-dependent fitness effects in a collection of beneficial mutations from the early generations of the *E. coli* LTEE. These mutations represent well-characterized targets of adaptation that were previously considered to have simple, unconditionally beneficial effects^46^. Using highly precise flow cytometry-based competition assays, we measure fitness effects across multiple frequencies and test fundamental assumptions about the constancy and transitivity of fitness relationships. We find that frequency-dependent fitness effects are ubiquitous, occurring in the majority of strain pairs tested. Through detailed analysis of within-growth cycle dynamics, we identify resource competition as a key mechanism underlying frequency-dependence and demonstrate that simple ecological interactions can generate complex fitness relationships, even in highly controlled laboratory environments.

The implications of widespread frequency-dependent fitness effects extend across multiple domains. If fitness effects typically depend on population composition rather than remaining constant, this would affect predictions about evolutionary dynamics, the interpretation of epistatic interactions, and the design of competition experiments to measure fitness effects. Moreover, the ubiquity of frequency-dependent effects could explain the persistence of genetic diversity in laboratory populations and provide insights into the mechanisms that generate and maintain the extraordinary diversity observed in natural microbial communities^47–51^.

## Results

### Beneficial mutations exhibit pervasive frequency-dependent fitness effects

In simple, ecologically unstructured microbial populations, the fitness effects of mutations are typically assumed to be constant properties, independent of frequency. We tested this assumption using one of the best-characterized sets of beneficial mutations in experimental evolution: the first five beneficial mutations to fix in one of the twelve LTEE populations (Ara-1), which are also common, parallel targets of adaptation across the LTEE more broadly^46^ (Figure 1a). Reconstructed individually and in all 32 (= 2^5^) combinations on the clean background of the LTEE ancestor REL606, the fitness effects of these mutations have been measured exhaustively to dissect epistasis, revealing a smooth, single-peaked adaptive landscape governed by diminishing-returns epistasis that helps account for the deceleration of adaptation over the experiment^45,46^. Throughout this body of work, these mutations have been modeled as having constant, unconditionally beneficial effects. The mutants are labeled: *G*, SNP in the *glmUS* promoter; *R*, deletion of the *rbs* operon; *P*, deletion of the *pykF* gene; *S*, SNP in the *spoT* gene; and *T*, SNP in the *topA* gene. Here, we consider all single and double mutants.

**Figure 1.**
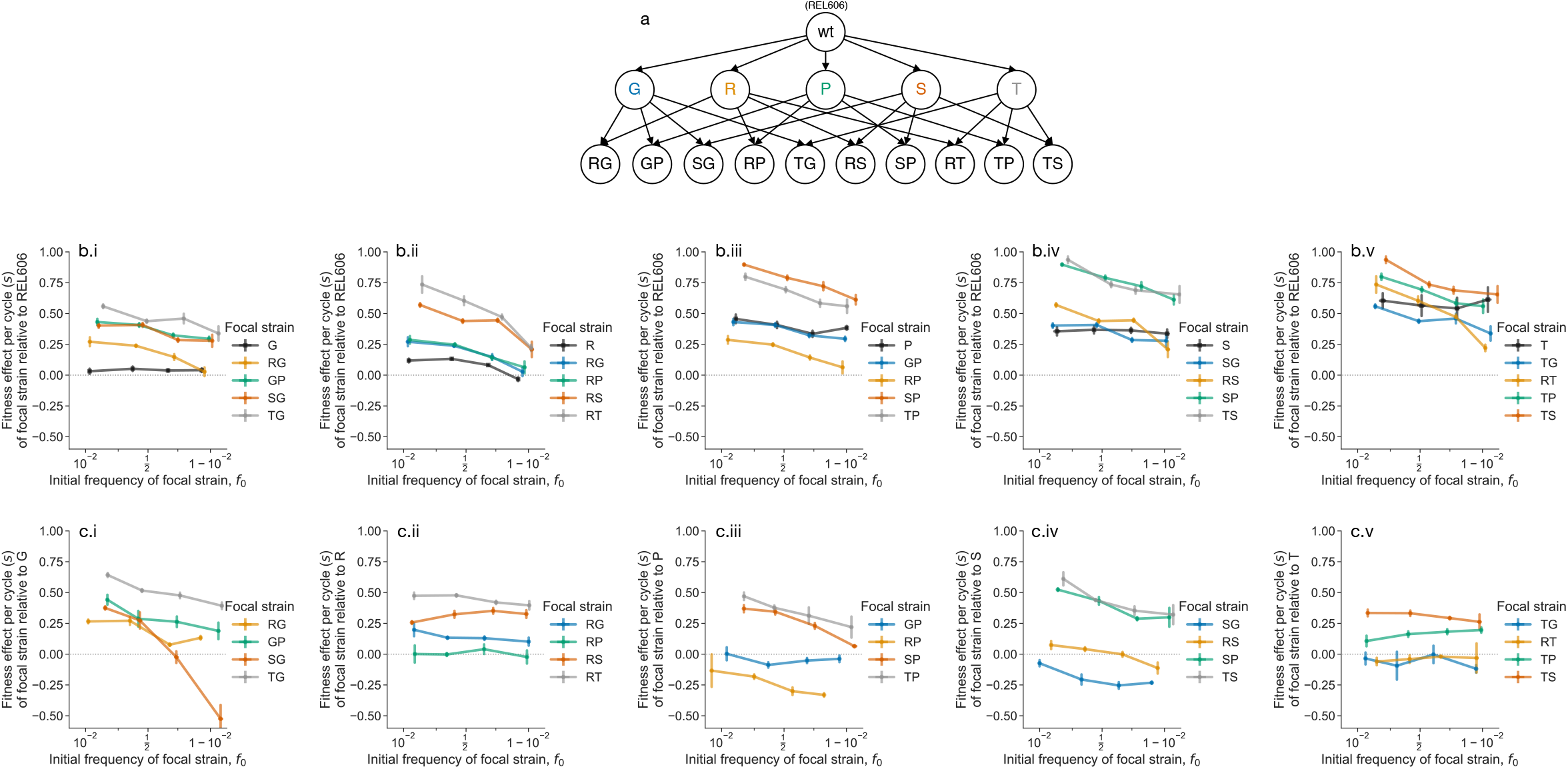
Ubiquitous frequency-dependent fitness effects. (**a**) We study a set of single and double mutants, all derived from the ancestor of the *E. coli* LTEE, REL606. Genotypes are denoted by nodes, and ancestral relationships denoted by arrows. (**b**) We measured the fitness effects, *s*, of all single and double mutants against REL606 as a function of frequency. Each subpanel (i-v) shows the measured frequency-dependent fitness effects for each single mutant and derived double mutants. Note that we plot the frequency-dependent fitness effects of double mutants twice–once on each subpanel corresponding to an immediate ancestral single mutant–to allow for comparison. (**c**) We additionally measured the frequency-dependent fitness effects of all double mutants against their single mutant ancestors. Error bars represent standard errors across biological replicates (*n* =4∼8) (see Table S1).

To measure fitness effects via flow cytometry, we tagged each mutant with either a red or yellow fluorescent protein (RFP/YFP); the fluorescent protein gene was integrated into a putatively neutral, intergenic genomic site via a Tn7 transposon system^52^. Consistent with previous work^27^, the choice of fluorescent protein had no measurable effect on fitness (FigureS4). The LTEE populations evolve in a simple daily dilution environment, where one one-hundredth of the population is transferred to fresh glucose minimal media every 24 hours. We initiated competitions in the LTEE environment at four different starting frequencies, around 1%, 20%, 80%, and 99% (approximately equally spaced on the logit axis). We measured frequencies ( *f* ) over two growth cycles, calculating the fitness effect (*s*) of the focal strain relative to the competitor, on a per-cycle basis, as the slope of the logit-transformed frequencies^53,54^:

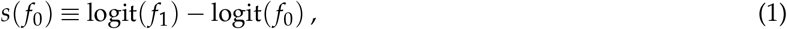

where 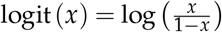. We also computed fitness as the ratio of Malthusian parameters,

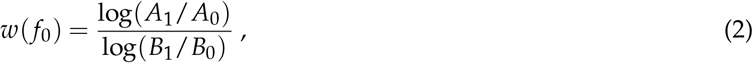

where *A*_*t*_ and *B*_*t*_ are the population sizes of the focal and competitor strains, respectively, at time *t*. Our results generally do not depend strongly on the choice of fitness measure, so we generally report fitness effects (*s*) in the main text and Malthusian fitness (*w*) in the supplement.

Contrary to prior expectations, we observed substantial frequency-dependent effects in most strain pairs (Figures 1b-c, S1). While most focal strains maintained positive mean fitness effects relative to the ancestral competitor REL606, the magnitude of these effects varied significantly with frequency. In at least two cases (SG vs. G and RS vs. S), the frequency-dependent fitness relationship crossed zero, indicating potential for stable coexistence (Figure 1c.i&.iv). At least three double mutants (RP, SG, TG) showed lower mean fitness than their single mutant ancestors in some conditions–a signature of sign epistasis, consistent with prior data^46^ (Figure 2a-b).

**Figure 2.**
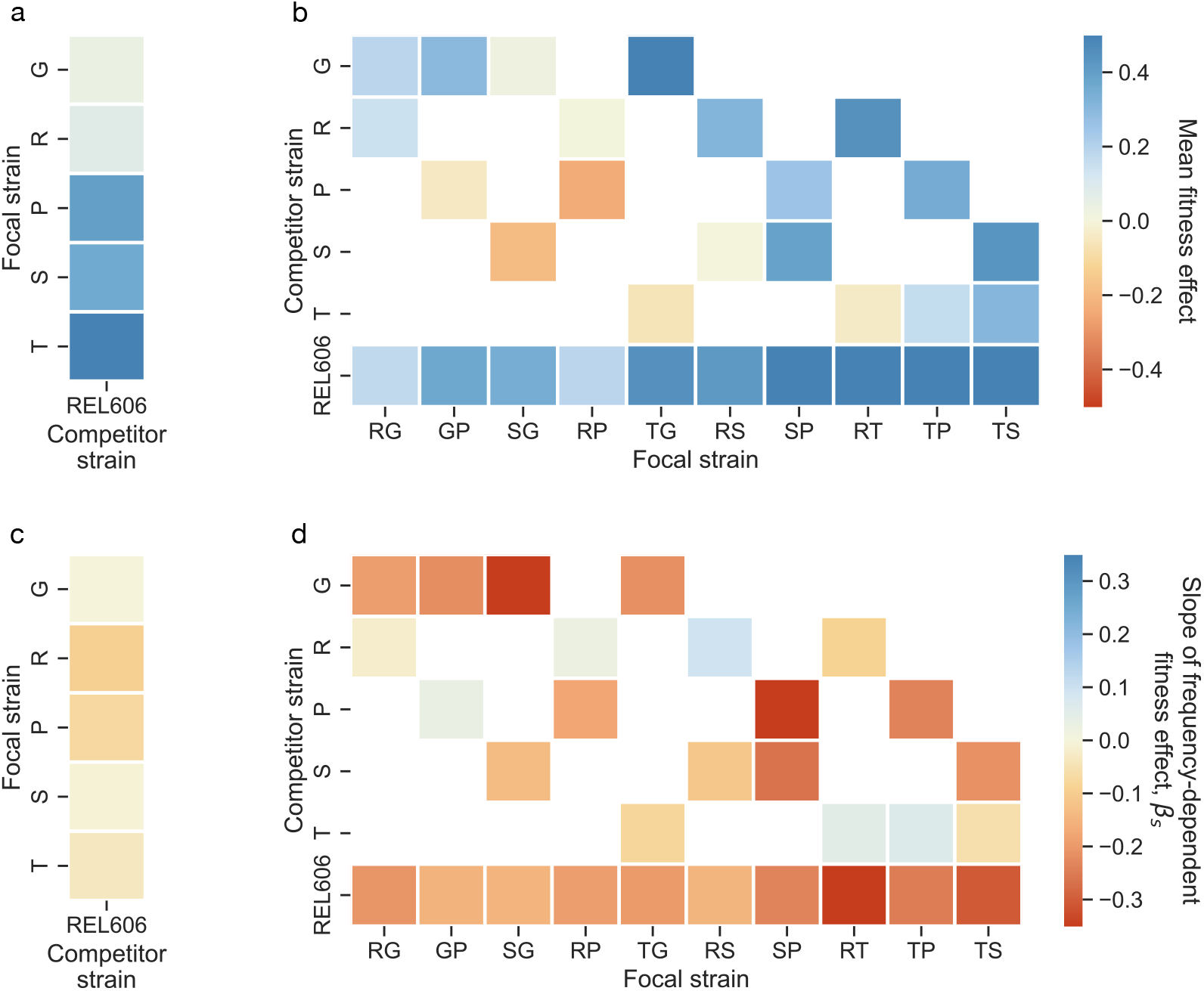
Summaries of frequency-dependent fitness effects. (**a-b**) Mean fitness effect of the focal strain relative to the competitor strain, across biological replicates and measured frequencies, of all competitions. (**c-d**) Mean slope of frequency-dependent fitness effects for all competitions, of the focal strain relative to the competitor strain. Slopes obtained through orthogonal distance regression (SI section S2.4).

Approximately 80% of competitions (at *p <* 0.01, FDR-corrected) exhibited clear negative frequency-dependence, where fitness effects declined as mutant frequency increased (Figures 2c-d, S8). This pattern would cause fixation dynamics to slow as beneficial mutations approach fixation, extending the time required for mutants to sweep through populations. Indeed, Wright-Fisher-like simulations (in the strong selection-weak mutation regime) reveal that the negative frequency-dependent fitness effects observed here typically increase the average mutant fixation time by hundreds of generations (Figure S7). Interestingly, two competitions (RS vs. R and TP vs. T) showed significant positive frequency-dependence. Double mutants exhibited frequency-dependent effects more frequently than single mutants–only three of five single mutants showing measurable frequency-dependence against REL606, compared to 27/30 competitions involving double mutants. The ubiquity of frequency-dependent effects across these well-characterized beneficial mutations reveals that ecological interactions are the norm rather than the exception, even in laboratory environments designed to minimize complexity.

### Predictable patterns emerge in frequency-dependent fitness effects

Given the widespread frequency-dependent fitness effects among the tested clonal strains, we sought to understand if there are any predictable factors that affect the magnitude of frequency-dependence. Consistent with widespread negative frequency-dependence, fitness effects where the focal strain is in the minority, *s*_*inv*_, are higher than when the focal strain is in the majority, *s*_*high*_, on average (Figure 3a). Moreover, the relationship between *s*_*inv*_ and *s*_*high*_ shows a slight curvature at higher absolute invasion fitness effects. This curvature is reflective of a general negative dependence of the slope (*β*_*s*_) of the frequency-dependence on *s*_*inv*_ (Figure 3b). This negative correlation is quite strong, with absolute correlation coefficients around 50-60%. We find similar patterns when considering Malthusian fitness (Figure S9).

**Figure 3.**
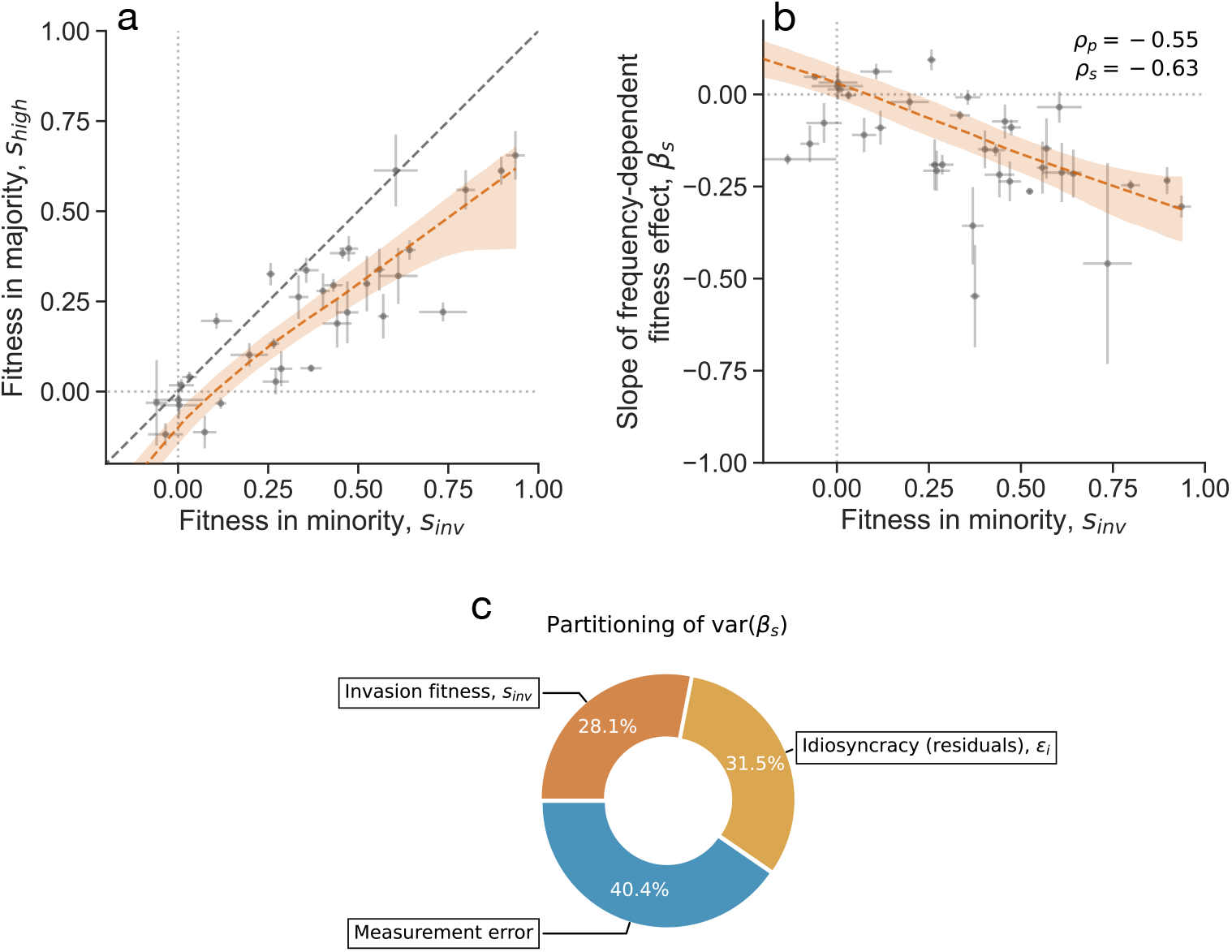
Statistical patterns of frequency-dependent fitness effects. (**a**) Comparison of low-frequency (*s*_*inv*_) with high-frequency (*s*_*high*_) fitness effects for all competitions. (**b**) The average slope of frequency-dependent fitness effects is negatively correlated with the fitness effect of the focal strain in the minority. *ρ*_*p*_ represents Pearson’s correlation; *ρ*_*s*_ represents Spearman’s correlation. The red lines represent a LOWESS fit. All error bars represent standard errors. (**c**) Partitioning the variance in frequency-dependent fitness effect slopes using ANCOVA into three components: invasion fitness, biological idiosyncrasy, and measurement error.

We fit ANCOVA models (Supplement section S2.5) to the data to partition the variance among competitions of the frequency-dependent fitness effect slope into three components: invasion fitness, biological idiosyncrasy, and measurement error. We find that measurement error can explain about 40% of the variance in *β*_*s*_, and biological idiosyncrasy (unexplained variance) and invasion fitness each explain another 30%. Thus, we are able to explain roughly half of the non-technical variation in *β*_*s*_ through invasion fitness measurements alone. The strong predictive relationship between invasion fitness and frequency-dependent slope suggests that these ecological interactions follow systematic, predictable rules.

### Fitness relationships violate transitivity assumptions

Evolutionary theory typically assumes that fitness effects are transitive and additive. Under this assumption, if we measure fitness effects *s*_*ik*_ of strain *i* relative to strain *k* and *s*_*jk*_ of strain *j* relative to strain *k*, we should be able to predict the fitness effect of strain *i* relative to strain *j, s*_*ij*_ = *s*_*ik*_ − *s*_*jk*_. Deviations from this relationship indicate that ecological interactions affect fitness in context-dependent ways. We refer to all such deviations as fitness non-transitivity, *ν* = *s*_*ij*_ − *s*_*ik*_ + *s*_*jk*_ (observed - expected fitness effect).

We tested transitivity by comparing predicted and observed fitness effects across strain triads, focusing on cases where one strain comprised the vast majority or minority of the population. We examined eight possible definitions of frequency-dependent non-transitivity (see Supplement sectionS2.2), presenting two natural cases here (Figure 4a-b; remaining cases in Figure S12).

**Figure 4.**
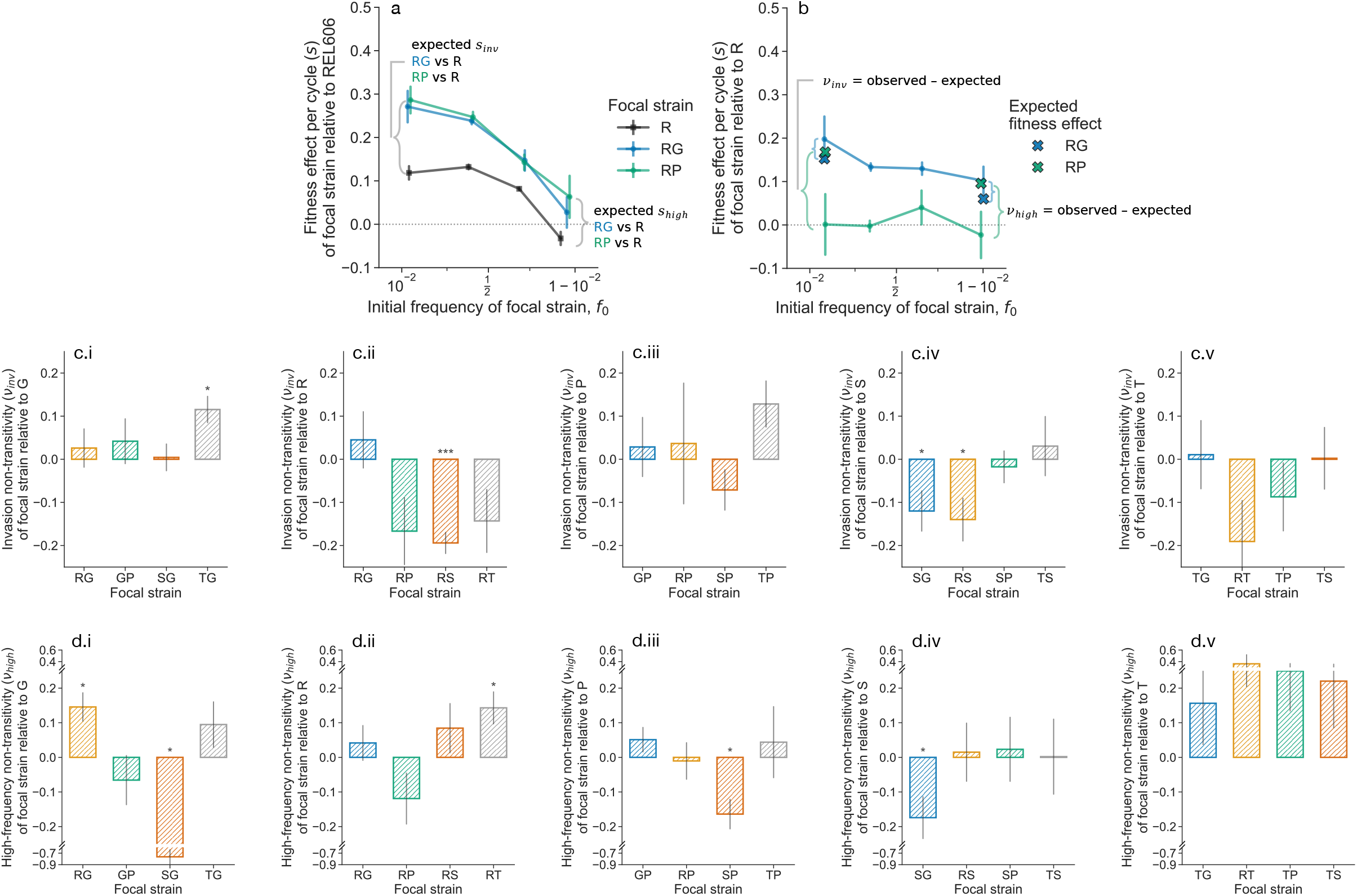
Non-transitivity in fitness effects. (**a-b**) Non-transitivity is defined as the deviation between expected fitness effects (if fitness effects between two competitions are additive) and observed fitness effects in a third experiment. Here, we show results from two definitions of non-transitivity at (**c**) low and (**d**) high frequencies; additional definitions are shown in Figure S12. Each column (i-v) represents non-transitivity of double mutants against different single mutants: (i) G, (ii) R, (iii) P, (iv) S, (v) T. Note that y-axis scales differ between rows. \**p <* 0.05, \*\**p <* 0.01, \*\*\**p <* 0.001, post-FDR correction.

**Figure 5.**
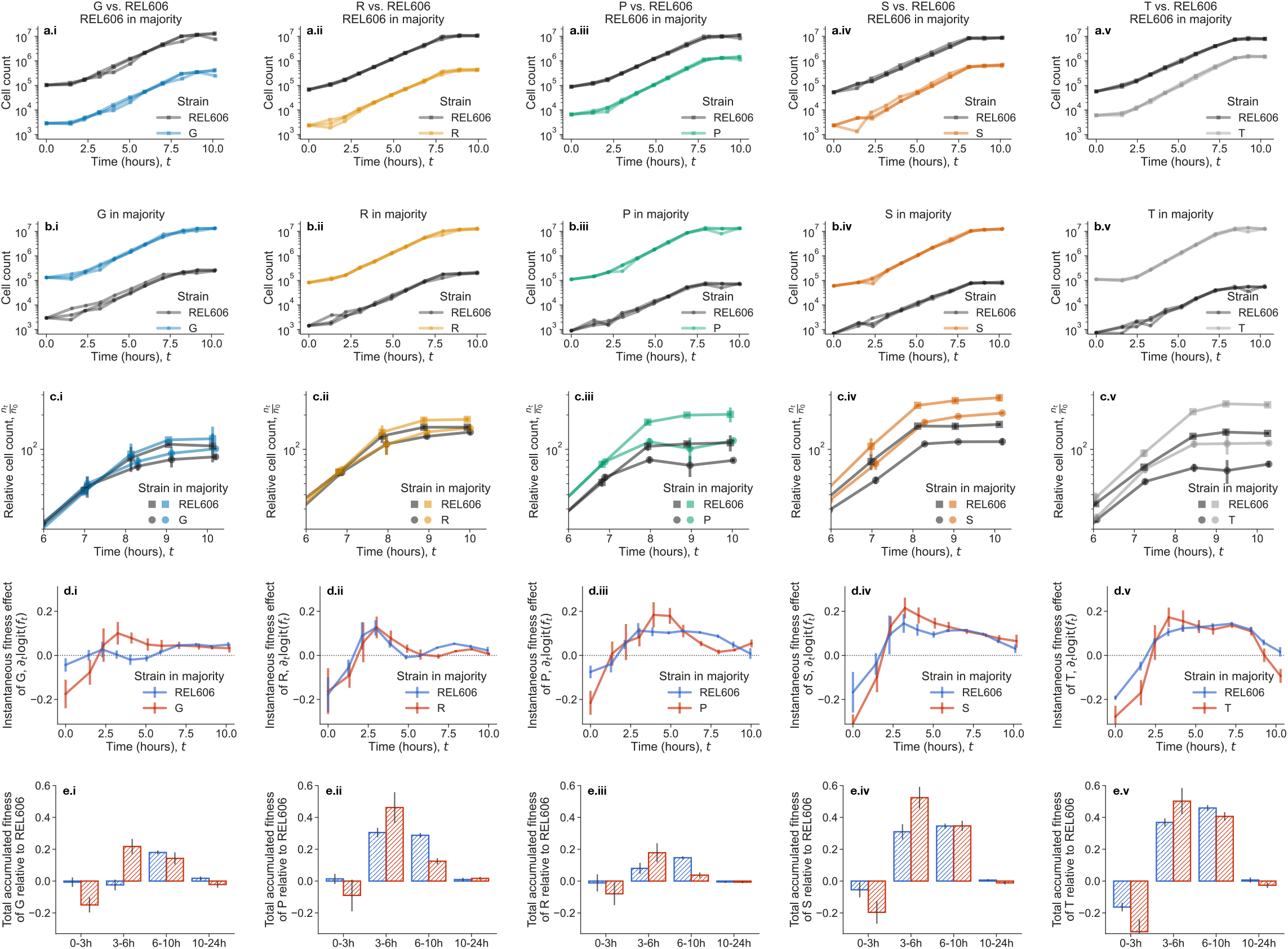
Within-growth cycle dynamics reveal ecological dynamics. (a-b) Dynamics of cell counts over the course of the first ∼10 hours of the growth cycle; where either (a) REL606 is in the majority of the population, or (b) the mutant is in the majority of the population. Trajectories are shown for all three biological replicates per condition. (c) Comparison of mean relative cell count trajectories (normalized to initial count) at the end of exponential phase. Generally, when the mutant is in the majority, all strains tend to exit exponential phase earlier than when REL606 is in the majority. (d) Estimates of the instantaneous, time-dependent fitness effects of the mutant, relative to REL606. (e) Total fitness accumulated over specified intervals of time, i.e. effectively integrating the curves in (d) over intervals in time. All error bars correspond to standard errors.

We observed clear and significant violations of transitivity in several strain combinations, where observed fitness non-transitivity, *ν*, typically ranged around *±*20%. Under four other definitions of non-transitivity, there are a higher number of significant cases of non-transitivity (Figure S12). These non-transitive effects indicate that fitness effects are context-dependent; they could substantially alter evolutionary dynamics by favoring or disfavoring mutations based on the composition of competing genotypes in the population.

### Niche construction drives frequency-dependent fitness effects

To understand the ecological basis of frequency-dependent fitness effects, we measured within-growth cycle dynamics for competitions between each single mutant and REL606 at both high and low frequencies (Figure 5a-b). We subsampled cocultures approximately hourly for ten hours, covering exponential phase and the transition to stationary phase, with an additional measurement at 24 hours. Population sizes are approximately constant from 10-24 hours, as expected (Figure S13).

We find that the frequency-dependence changes substantially within the growth cycle, with the net per-cycle effect reflecting the balance of opposing dynamics at different growth phases. The “instantaneous” fitness effects changed dramatically over time for each competition (Figure 5d). Even mutants (G, S, T) that showed no significant overall frequency-dependence exhibited transient frequency-dependent advantages and disadvantages that quantitatively canceled out over the full growth cycle (Figure S15). Competitions generally showed three qualitatively consistent phases (Figure 5e): (1) mutants initially grew slower than REL606, but performed relatively better when REL606 was in the majority; (2) mutants gained fitness advantages around 3-6 hours, with stronger advantages when the mutant was in the majority; and (3) around 6-10 hours, mutants generally performed better with REL606 in the majority. These frequency-dependent differences in fitness trajectories suggest that ancestral and mutant strains modify their shared environment in different ways; these environmental modifications appear strong enough to noticeably influence fitness trajectories.

We can further investigate the causes of transient fitness dynamics by examining population growth rates over time. Fitness effects and growth rates are related in a straightforward way: the difference in growth rates between mutant and ancestor equals the fitness effect, ∂_*t*_ logit( *f* ) = −∂_*t*_ log(*n*_*mut*_) ∂_*t*_ log(*n*_*wt*_). Growth rates for all strains follow an inverted-U trajectory across all conditions (Figure S14). This pattern is expected for bacterial growth dynamics–growth rates typically peak in mid-exponential phase before declining toward stationary phase. However, we observed substantial quantitative differences in growth trajectories between frequency conditions. Both REL606 and mutant growth rates depend on population composition, with both strain types achieving higher growth rates when REL606 dominates, particularly at the beginning and end of exponential phase.

From our within-cycle growth curve data, we find that frequency-dependence during late-exponential phase typically accounts for 10–40% of the absolute per-cycle fitness effect, depending on the strains involved (Figure S15). This late-exponential frequency-dependence has a straightforward explanation: raw growth curves (Figures 5c, S20) show that populations enter stationary phase earlier when mutants dominate, meaning that the duration of exponential growth itself depends on population composition. To investigate the mechanistic basis of this pattern, we analyzed a simple consumer-resource model (Supplement Section S4). We assume that wild-type and mutant strains grow by consuming a single, exhaustible resource, with strain-specific growth rates (*r*_*wt*_, *r*_*mut*_) and resource conversion factors (*a*_*wt*_, *a*_*mut*_). Analyzing the dynamics in the high-and low-frequency limits, we show that resource depletion time *T* scales inversely with the growth rate of the dominant strain, and is thus frequency-dependent: 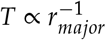. This frequency-dependent saturation time has been previously analyzed in theoretical models of serial dilution cultures^55,56^, where selection is shown to depend on both growth rate and lag time differences accumulated over a growth duration that itself depends on the competing strains’ traits. Importantly, this effect induces a frequency-dependence in the absolute fitness difference per cycle, *s* = *T*(*r*_*mut*_ − *r*_*wt*_), because *T* varies with frequency. However, it does not generate frequency-dependence in the ratio of Malthusian parameters (relative fitness *w*), which is a dimensionless quantity independent of *T*. Thus, frequency-dependent *T* changes how much absolute fitness difference accumulates per cycle, but does not, on its own, alter the competitive ranking of strains. That being said, the frequency-dependent *s* still can have strong evolutionary consequences, for example, by slowing the fixation of beneficial mutants (Figure S7). Overall, this provides a general mechanism for frequency-dependent fitness during late exponential phase: faster-growing mutants deplete resources more quickly when dominant, shortening the time available for fitness advantages to accumulate.

We can estimate the contribution of resource depletion to frequency-dependent fitness by comparing fitness effects (*s*) with Malthusian fitness (*w*). Fitness effects have units of inverse time, but Malthusian fitness is a dimensionless ratio and therefore does not depend on the frequency-dependent depletion time *T*. Any frequency-dependence in *w* must therefore arise through other mechanisms. By comparing the frequency-dependent slopes of *s* and *w* across all competitions (Supplement Section S4.3), we estimate that resource depletion dynamics account for 10–50% of frequency-dependent fitness, depending on the strain pair (Figure S16). Thus, the total frequency-dependence is partially driven by the simple consequences of nutrient limitation, but is generally primarily driven by other, possibly idiosyncratic, ecological processes. These results demonstrate that subtle ecological interactions between closely related genotypes occur even in simple, controlled, and “constant” laboratory environments.

## Discussion

Our demonstration that frequency-dependent fitness effects are ubiquitous among beneficial mutations from the early LTEE challenges assumptions about the typicality of constant fitness effects in microbial evolution^1,46^. We find that even in highly controlled laboratory environments, subtle ecological interactions generate complex evolutionary dynamics that most models of microbial evolution do not account for.

Perhaps the most surprising aspect of our results is how common frequency-dependent fitness effects are. Rather than unusual cases requiring special explanation, frequency-dependence appears to be the norm. This is particularly striking given that our study system–the LTEE environment, with its simple glucose-limited medium and controlled conditions–was specifically designed to minimize ecological complexity^1,31^. The prevalence of frequency-dependence may explain why coexistence has arisen so predictably in the LTEE^43^: if negative frequency-dependent fitness effects that cross zero are common, establishing coexistence may be relatively easy, provided further adaptation does not eliminate the nascent ecosystem. Recent theory for diverse ecological communities likewise predicts that feedbacks arising as mutants increase in abundance can modify the relationship between invasion fitness and subsequent dynamics^57^, suppress the fixation of moderately beneficial mutations, and prolong parent–mutant coexistence^58^. Frequency-dependent fitness effects have also recently been reported in experimental *Saccharomyces cerevisiae* cultures^59^, suggesting these dynamics may be widespread across microbial systems.

It is worth emphasizing that the LTEE, and our assays, use daily batch transfers rather than continuous culture. Batch dynamics impose a natural cycle of feast and famine, with successive temporal niches (lag, exponential, transition, stationary) over which strains can differ^55,56^. Such temporal niche structure has been argued to be sufficient to seed ecological diversification from physiological trade-offs^60^. In a chemostat or other continuous-flow environment, where a single steady-state niche dominates, we may expect the prevalence and magnitude of frequency-dependence among closely related single and double mutants to be smaller. However, the presence of more than one temporal, resource, or other niches would open back up the possibility of significant frequency-dependent fitness effects.

The within-cycle structure of frequency-dependence–transient costs during lag and early exponential phase (0-3 h), advantages during mid-exponential phase (3-6 h), and partial cancellation in late exponential phase (6-10 h)– points toward possible physiological mechanisms that may differ across mutants. The 0-3 h interval corresponds to lag and early exponential growth, where strains must remodel their proteome upon resuming growth from stationary phase; mutations that increase peak growth rate often do so by reallocating proteome to ribosomes or central metabolism^61,62^, which can lengthen lag or slow the initial exit from stationary phase–a classic growth-rate-vs-lag trade-off documented in *E. coli*^55,60,63^. The mid-exponential advantage (3-6 h) is the regime in which steady-state growth rate dominates. The late-exponential (6-10 h) is dominated by frequency-dependent resource depletion times mediated by simple resource competition dynamics: when faster-growing genotypes dominate, they deplete resources more quickly, reducing the time for fitness differences to accumulate.

Although strain-to-strain variability in phase-dependence likely reflects the different physiological strategies by which each mutation increases growth rate, all five single mutants nonetheless share a consistent relative fitness deficit during lag and early exponential growth, suggesting that a lag-associated cost may be a common pleiotropic consequence of early growth-rate adaptation. This pattern stands in apparent tension with earlier work showing that LTEE populations evolved shorter lag phases over the first 2,000 generations^64^. We hypothesize that this contrast reflects a two-stage process: relatively large-effect mutations are substituted early in adaptation^65^, increasing exponential growth rate while imposing deleterious pleiotropic effects on other growth-cycle traits, putatively mediated through the growth-rate-lag-time trade-off; subsequent compensatory substitutions then repair these costs while preserving the initial growth-rate benefit^66,67^. Under this view, the individually reconstructed mutations expose the pleiotropic costs of early adaptive steps, whereas later substitutions or epistatic combinations among them could produce the shorter-lag phenotype observed in evolved populations, which would imply that the growth-rate-lag-time trade-off is not immutable but can be modified by adaptive evolution.

Ultimately, our results suggest that the apparent simplicity of laboratory evolution experiments may be deceiving. Even in the most controlled environments, evolution operates through complex ecological interactions that deform fitness landscapes, determine competitive outcomes, and influence the maintenance of genetic diversity. Rather than simple hill-climbing on static fitness landscapes, microbial communities navigate dynamic fitness spaces shaped by ecological interactions, epistatic constraints, and environmental feedback. Recognizing and incorporating these interactions into our theoretical and experimental frameworks will be essential for developing a more complete understanding of evolutionary processes across all scales of biological organization.

## Methods

See supplementary information.

## Acknowledgements

We extend a special thank you to Richard Lenski, for providing us with especially helpful feedback on an earlier version of this manuscript. We thank Adam Arkin, Michael Desai, and all members of the Hallatschek lab (past and present) for helpful discussions and advice on the project. We thank Tim Cooper, Richard Lenski, and Vaughn Cooper for sending us the LTEE-derived strains and populations, along with experimental advice and feedback. This work was supported by the National Institute of General Medical Sciences of the NIH under award R01GM149827 and by a Humboldt Professorship of the Alexander von Humboldt Foundation. JAA acknowledges support from an NSF graduate research fellowship, a Berkeley fellowship (from UC Berkeley), and Lloyd and Brodie scholarships (from UC Berkeley Dept of Bioengineering).

## Supplementary Information

### S1 Experimental methods

#### S1.1 Growth conditions, media and strains

All experiments described here were conducted in Davis Minimal (DM) base medium, composed of 5.36 g/L potassium phosphate (dibasic), 2 g/L potassium phosphate (monobasic), 1 g/L ammonium sulfate, 0.5 g/L sodium citrate, 0.01% magnesium sulfate, and 0.0002% thiamine HCl. The specific medium used in both the Long-Term Evolution Experiment (LTEE) and all assays presented here was DM25–i.e. DM supplemented with 25 mg/L glucose.

For coculture experiments, we began by (i) inoculating the desired strain into 1 mL of LB supplemented with 0.2% glucose and 20 mM pyruvate directly from glycerol stock. (ii) Following overnight incubation, cultures were washed three times in DM0 (DM lacking a carbon source) by centrifugation at 2500×*g* for 3 minutes, removal of the supernatant, and resuspension in DM0. (iii) The washed culture was then diluted 1:1000 into 1 mL of DM25; in general, we grew 1 mL cultures in glass 96-well plates (Thomas Scientific 6977B05). (iv) Cultures were incubated at 37°C for 24 hours in a shaking incubator. (v) The following day, cultures were transferred 1:100 into fresh DM25 and incubated under identical conditions. (vi) After another 24 hours of growth, we mixed selected cultures at defined frequencies and diluted the mixtures 1:100 into fresh DM25. (vii) Cultures were incubated again for 24 hours under the same conditions, after which experimental measurements were initiated.

The daily transfers and fitness measurements of the *E. coli* Long Term Evolution Experiment (LTEE) were generally performed in 50mL glass flasks filled with 10mL DM25. We performed a set of control experiments in those flasks to investigate if there were significant fitness differences when competitions were performed in flasks versus glass 96 well plates. For the flask control experiments, we prepared cultures in the same manner as with previous experiments, following steps (i-ii). We modified step (iii) by diluting the washed culture 1:1000 into 10mL of DM25 in a 50mL glass flask. The remaining steps (iv-vii) were performed as previously described, with the exception that the culture was transferred 1:100 into 10mL DM25 in a 50mL glass flask. We found that the fitness measurements in flasks were in quantitative agreement with the fitness measurements in the glass 96 well plates (Figure S5).

#### S1.2 Fluorescent tagging

Strains carried fluorescent protein markers integrated at the *attTn7* locus via a miniTn7 transposon system, as previously described [1, 2]. Each strain either expressed sYFP2 or mScarlet-I. Briefly, we transformed a conjugative *E. coli* donor strain (MFDpir; DAP auxotroph) with a temperature-sensitive plasmid that expresses the miniTn7 machinery, with a chloramphenicol resistance gene and either sYFP2 or mScarlet-I between right and left Tn7 recongition sequences. We then conjugated the donor with the strain of interest by growing the strains together on solid media (LB/agar supplemented with 0.2% glucose and 0.3 mM DAP) for about 24 hours at 30°C. After growth, we scraped up the resulting lawn, washed three times in DM0, and streaked the culture on DM2000/agar plates supplemented with 20 µg/mL chloramphenicol. After 48 hours of growth at 37°C, we picked several candidate transconjugant colonies, testing for successful integration via PCR, and for successful loss of the plasmid by ability to grow on LB/ampicillin media (plasmid backbone has an ampicillin resistance gene). Successful clones were grown in DM2000 and saved as glycerol stocks.

#### S1.3 Flow cytometry

All population-level measurements were performed using a ThermoFisher Attune Flow Cytometer (2017 model) located at the UC Berkeley QB3 Cell and Tissue Analysis Facility (CTAF). Samples were loaded into a round-bottom 96-well plate compatible with the autosampler. For each run, the instrument performed one wash and mixing cycle before measurement. To prevent cross-contamination, 50µL of bleach was run through the autosampler between samples.

Fluorescence detection was performed as follows: sYFP2 using the BL1 channel (488 nm laser, 530/30 nm bandpass filter); and mScarlet-I using the YL2 channel (561 nm laser, 620/15 nm bandpass filter). Cell counts and strain frequencies were extracted from raw data using a previously validated analysis pipeline [1]. We applied a previously described gating strategy [1], using threshold-based gates to distinguish fluorescent events from debris and background noise (see Figure S3).

### S2 Frequency-dependent fitness measurements

To measure frequency-dependent fitness effects of different pairs of clonal strains, we first prepared cultures as described in section S1.1. For each pair of strains, one was tagged with YFP, and the other RFP. We mixed each pair of strains at four volumetric fractions–0.01, 0.2, 0.8, and 0.99 (approximately equally spaced in logit space). We measured the relative frequencies of each strain at the end of a growth cycle via flow cytometry (day 0), then we propagated the cultures as usual (1:100 dilution into DM25). At the end of the second growth cycle, we measured the relative frequencies once more using flow cytometry (day 1). We generally used between 4 and 6 independent biological replicates for each measurement.

#### S2.1 Quantifying fitness effects

From the frequency measurements of clonal strains at days 0 and 1, we sought to quantify the relative fitness effects for each experiment. We denote 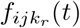 as the measured frequency of the focal strain *i*, relative to strain *j*, for experiment *k*_*r*_ at cycle number *t*. As previously mentioned, for any given set of experimental conditions (two strains and initial frequency), we will generally have 4-6 biological replicates; we denote *k* as a given experiment with a set initial frequency, and *r* as a given biological replicate. Then, following the standard population genetic definition of fitness effects, we calculate the fitness effect in a given experiment as,

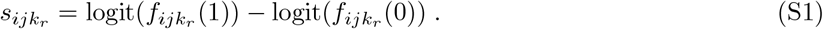

We noticed that the calculated 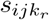 of a few biological replicates appeared to be significantly different from the remaining replicates. It appeared that most of these instances were due to apparent errors in the flow cytometer, e.g. early termination, fluidics errors, bubbles, etc. Thus, we excluded outliers from further analyis. We defined an outlier as any 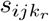 that is more than three standard deviations away from the median between all biological replicates, where standard deviation is approximated by the median absolute deviation. For normally distributed data, the relationship between standard deviation and median absolute deviation is, 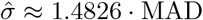, thus we use,

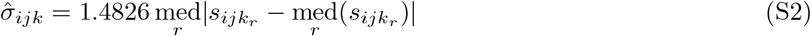

After outlier filtering, we computed our final estimates of *s*_*ijk*_ by simply averaging over 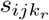 for all biological replicates *r*. We obtained standard errors for *s*_*ijk*_ and the initial frequency *f*_*ijk*_(0) by using the sample standard deviation.

#### S2.2 Quantifying non-transitivity

Under typical null models used in evolutionary biology, fitness effects are generally assumed to be additive and transitive. This arises from an assumption that the fitness of a clone is independent of the pairwise competition used to measure fitness effects. To see this, we note that the fitness effect, *s*_*ij*_(*E*), in environment *E*, of a given clone *i* relative to a competitor *j* can generically be decomposed into the difference in *fitness, x*(*E*), of each clone,

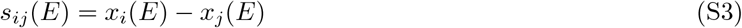

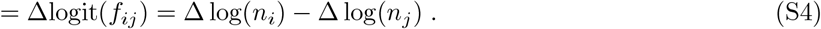

We denote *f*_*ij*_ as the frequency of *i* relative to *j*, and *n*_*i*_ as the population size of *i*. The environment *E* might be set either by any ecological interactions, or abiotic factors. If fitnesses are constant across competitions/environments, then if we measure fitness effects of a triplet of clones, we should recover a simple relationship between the measurements,

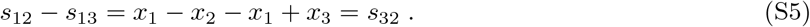

However, in general, this relationship will not hold if fitnesses depend on the specific ecological interactions of a given competition. We can quantify the deviation from transitivity (non-transitivity) between competition triplets with a quantity *ν*,

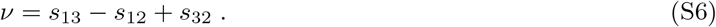

Of course, under this definition, *ν* = 0 when none of the three clones have competition-dependent fitnesses.

However, in the case that fitness effects are frequency-dependent, we must choose frequencies to use for fitness effect comparisons. We choose to focus on the cases where one strain is in the vast majority or minority of the population, i.e. when one of the frequencies tends to zero. We reasoned that this regime would allow for simplified interpretation of non-transitivity, as the environment will be dominated by the effects of the strain in the majority. Here, we denote *E*_*i*_ as the environment that is induced when strain *i* is in the vast majority of the population, *f*_*i*_ → 1. Under this constraint, there are eight possible different definitions of frequency-dependent non-transitivity. Two of these definitions appear especially natural (which we highlight in the main text): the case in which all environments are dominated by the competitor strains, comparing solely invasion fitness effects,

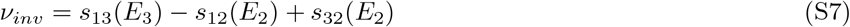

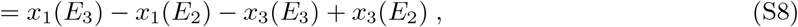

and the case in which all environments are dominated by the focal strains, comparing solely high-frequency fitness effects,

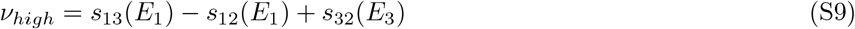

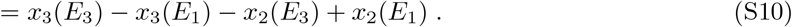

We see that the fitness of one strain cancels out in each definition, and we’re left comparing the fitness of two different strains in two different environments. In the case of quantifying the non-transitivity in our experimental data, we assign the labels: 1=double mutant; 2=wild-type (REL606); 3=single mutant. We show the results of such quantification in Figure 2. There are four other ways to construct defintions of non-transitivity so that the fitness of one strain gets canceled out,

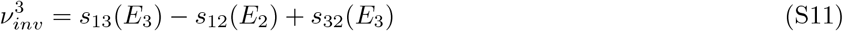

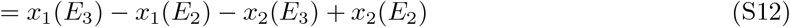

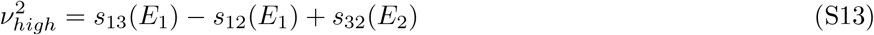

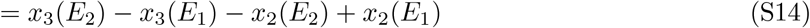

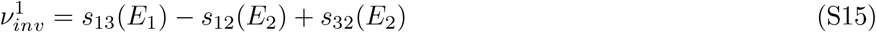

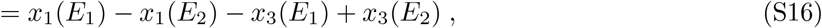

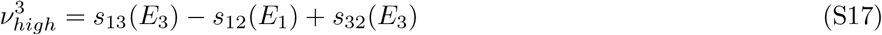

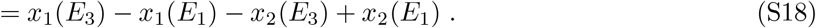

We show the results of quantifying non-transitivity under these definitions in Figures **xx and xx**.There are two additional ways to construct definitions of frequency-dependent non-transitivity such that none of the fitnesses get canceled out; these two definitions are simply linear combinations of prior definitions.

We sought to quantify not only the point estimates of *ν*, but also estimates of uncertainty. We computed the sample standard error of *ν* from the standard errors of the fitness effects 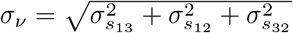. Then, we sought to compute p-values, testing the null hypothesis that *ν* = 0. We used a one-tailed, one-sample t-test, where we combined the degrees of freedom from all fitness effects using the standard Welch–Satterthwaite equation,

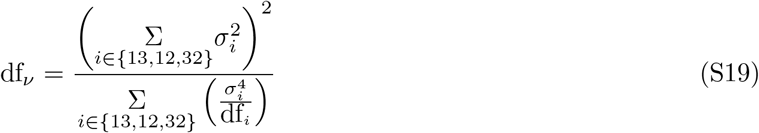

We used a standard Benjamini-Hochberg FDR correction to correct the p-values for multiple testing.

#### S2.3 Frequency-dependent epistasis

We wish to quantify additive and epistatic coefficients of double and single mutants. Our situation differs from the usual case, as we must compare fitness effects across different environments. For simplicity, we focus on fitness effects at invasion and high frequencies.

For any given environment *E* where a given clonal strain is in the majority, the additive coefficient of a given single mutant *i*, we simple call the additive coefficient the measured fitness effect *s*_*i*_(*E*). Then, to quantify the epistasis between two mutations *i* and *j*, we simply take the difference between the observed fitness effect of the double mutant from the expected value under additivity, *s*_*ij*_(*E*) − *s*_*i*_(*E*) − *s*_*j*_(*E*). Note that we drop the factor of two that appears in some models of epistasis. We calculate p-values as in the previous section, through standard one-sample t-tests. We present the results of this analysis in Figure S6.

#### S2.4 Quantifying frequency-dependent slopes and curvature

We wished to estimate the average slope (*β*_*s,i*_) and curvature (*y*_*i*_) of the frequency-dependent fitness effects (*s*(*f*_0_)) that we measured, for each set of competitions *i*. We computed a similar slope (*β*_*w,i*_) for frequency-dependent Malthusian fitness (*w*(*f*_0_)). Both frequencies and fitness effects have experimental noise associated with each measurement, which we wished to incorporate into our estimates of *β*_*s,i*_ and *y*_*i*_. We assume gaussian errors on both the measured frequencies, 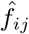, and measured fitness effects, *ŝ*_*ij*_, for each frequency measurement *j*. We took the measurement errors as the empirical standard errors, estimated from biological replicates. To estimate the slopes, we modeled the frequencies and fitness effects as,

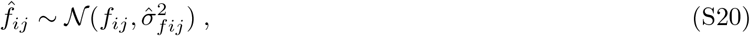

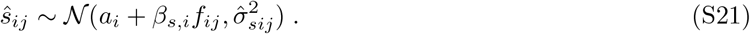

We used orthogonal distance regression to fit the slopes and intercepts, independently for each competition *i*, using the implementation in scipy (`scipy.odr`). Results are shown in table S1.

We also obtain an estimate of the error on the slope, 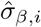. Analogously, to estimate the curvatures *y*_*i*_, we modeled the frequencies and fitness effects as,

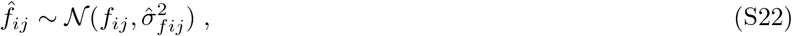

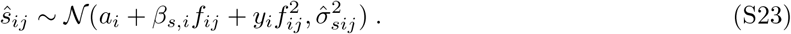

We performed the same procedure to infer the slopes, with the exception of excluding the points corresponding to the invasion fitness effects (Figure S10).

#### S2.5 ANCOVA model

Here we develop an ANCOVA model to model the slope of frequency-dependent fitness effects, *β*_*s,i*_. We consider three models: (1) a model that only considers measurement error and biological idiosyncrasy (i.e. unexplained variance/residuals), (2) a model that adds the invasion fitness, *s*_*inv,i*_, as a covariate, and (3) a model that adds the high-frequency fitness, *s*_*high,i*_, as a covariate. We only consider competitions where the focal strain has a higher number of mutations than the competitor strain.

For all of the models detailed below, we found the associated likelihoods to compute a maximum likelihood estimate of all parameters. We summarize the results from all of the models in Table S2.

##### Model 1

We begin by modeling the slope of frequency-dependent fitness effects for each pairwise competition *i, β*_*s,i*_, as linear combination of three components,

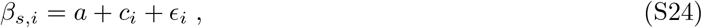

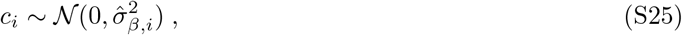

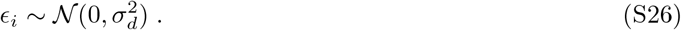

Here, *c*_*i*_ the measurement error, *ϵ*_*i*_ is unexplained variance, and *a* is the offset. We model *c*_*i*_ as a normal random variable with a known standard deviation, as the sample sizes used to estimate *β*_*s,i*_ are generally large enough such that the errors should be approximately normal. The likelihood for this model, combining all measurements, is

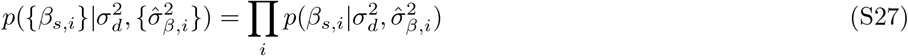

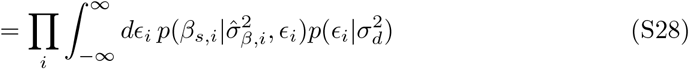

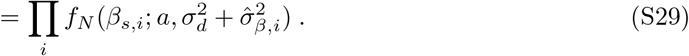

Here we denote *f*_*N*_ (*x*; *µ, σ*^2^) as a normal distribution of a random variable *x* with mean *µ* and variance *σ*^2^.

**Table S1:**
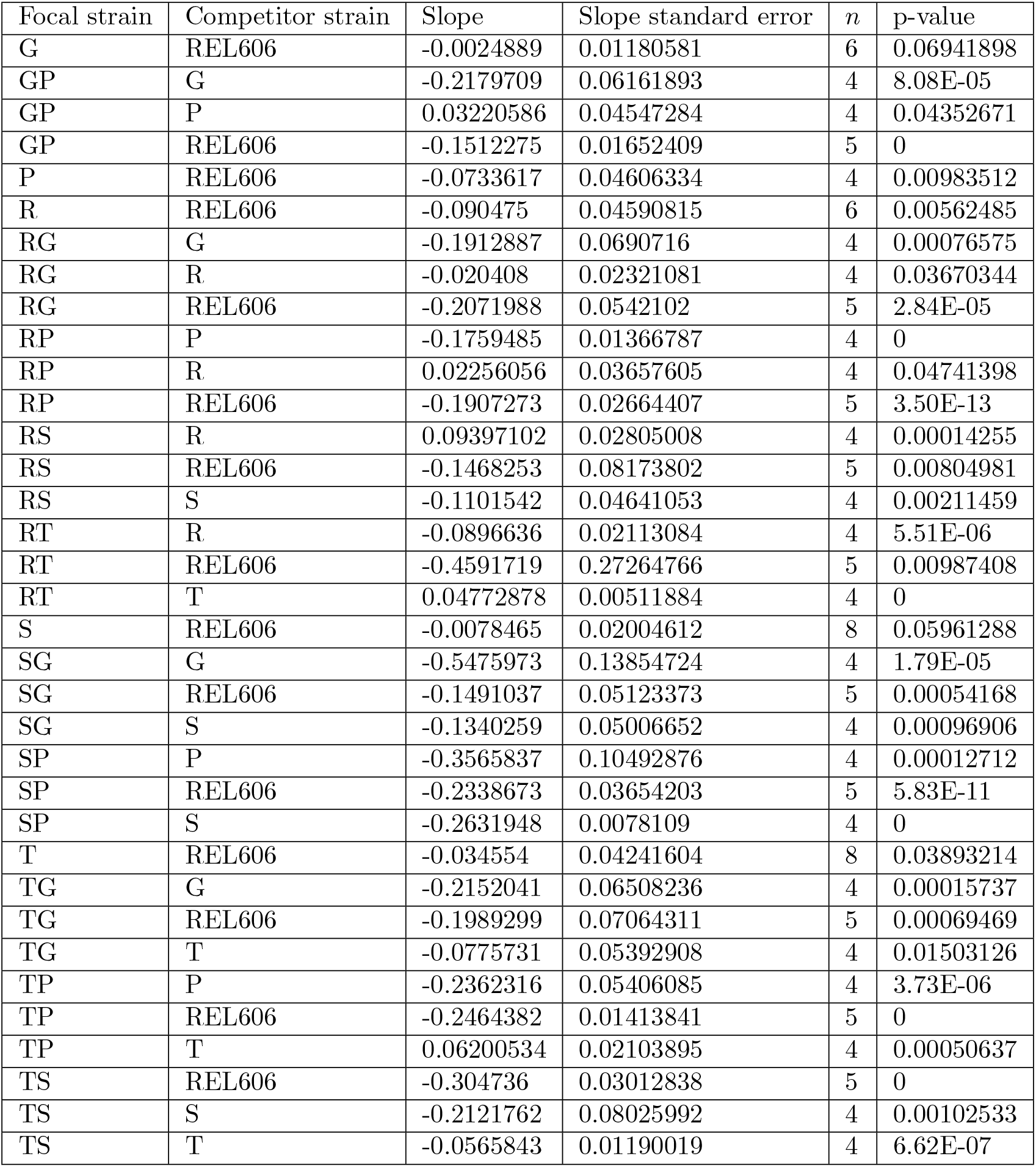
Frequency-dependent slopes for each competition. *n* is the number of biological replicates for a given competition. p-values are computed with a null hypothesis that the slopes are zero, and are corrected with a standard Benjamini-Hochberg FDR correction.

| Focal strain | Competitor strain | Slope | Slope standard error | $n$ | p-value |
| --- | --- | --- | --- | --- | --- |
| G | REL606 | -0.0024889 | 0.01180581 | 6 | 0.06941898 |
| GP | G | -0.2179709 | 0.06161893 | 4 | 8.08E-05 |
| GP | P | 0.03220586 | 0.04547284 | 4 | 0.04352671 |
| GP | REL606 | -0.1512275 | 0.01652409 | 5 | 0 |
| P | REL606 | -0.0733617 | 0.04606334 | 4 | 0.00983512 |
| R | REL606 | -0.090475 | 0.04590815 | 6 | 0.00562485 |
| RG | G | -0.1912887 | 0.0690716 | 4 | 0.00076575 |
| RG | R | -0.020408 | 0.02321081 | 4 | 0.03670344 |
| RG | REL606 | -0.2071988 | 0.0542102 | 5 | 2.84E-05 |
| RP | P | -0.1759485 | 0.01366787 | 4 | 0 |
| RP | R | 0.02256056 | 0.03657605 | 4 | 0.04741398 |
| RP | REL606 | -0.1907273 | 0.02664407 | 5 | 3.50E-13 |
| RS | R | 0.09397102 | 0.02805008 | 4 | 0.00014255 |
| RS | REL606 | -0.1468253 | 0.08173802 | 5 | 0.00804981 |
| RS | S | -0.1101542 | 0.04641053 | 4 | 0.00211459 |
| RT | R | -0.0896636 | 0.02113084 | 4 | 5.51E-06 |
| RT | REL606 | -0.4591719 | 0.27264766 | 5 | 0.00987408 |
| RT | T | 0.04772878 | 0.00511884 | 4 | 0 |
| S | REL606 | -0.0078465 | 0.02004612 | 8 | 0.05961288 |
| SG | G | -0.5475973 | 0.13854724 | 4 | 1.79E-05 |
| SG | REL606 | -0.1491037 | 0.05123373 | 5 | 0.00054168 |
| SG | S | -0.1340259 | 0.05006652 | 4 | 0.00096906 |
| SP | P | -0.3565837 | 0.10492876 | 4 | 0.00012712 |
| SP | REL606 | -0.2338673 | 0.03654203 | 5 | 5.83E-11 |
| SP | S | -0.2631948 | 0.0078109 | 4 | 0 |
| T | REL606 | -0.034554 | 0.04241604 | 8 | 0.03893214 |
| TG | G | -0.2152041 | 0.06508236 | 4 | 0.00015737 |
| TG | REL606 | -0.1989299 | 0.07064311 | 5 | 0.00069469 |
| TG | T | -0.0775731 | 0.05392908 | 4 | 0.01503126 |
| TP | P | -0.2362316 | 0.05406085 | 4 | 3.73E-06 |
| TP | REL606 | -0.2464382 | 0.01413841 | 5 | 0 |
| TP | T | 0.06200534 | 0.02103895 | 4 | 0.00050637 |
| TS | REL606 | -0.304736 | 0.03012838 | 5 | 0 |
| TS | S | -0.2121762 | 0.08025992 | 4 | 0.00102533 |
| TS | T | -0.0565843 | 0.01190019 | 4 | 6.62E-07 |

##### Model 2

We now extend our model of *β*_*s,i*_ to include the invasion fitness, *s*_*inv,i*_, as a covariate,

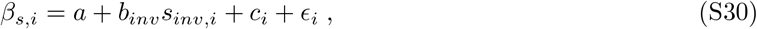

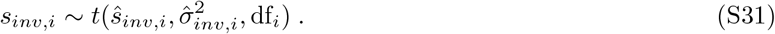

Here, *ŝ*_*inv,i*_ is the sample mean invasion fitness of competition *i*, 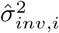 is the sample variance of the invasion fitness, and df_*i*_ = *n*_*i*_ − 1 are the degrees of freedom, related to the number of biological replicates for that measurement. We model *s*_*inv,i*_ as a non-centered, scaled *t*-distributed random variable, as we assume that underlying measurement error distribution is gaussian, but *n*_*i*_ is typically small, so we must account for errors in the estimated sample variance. Specifically, we use a probability distribution function of the form,

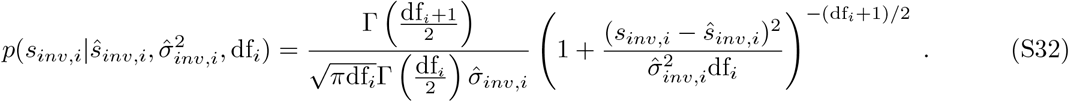

The full likelihood of the model is then

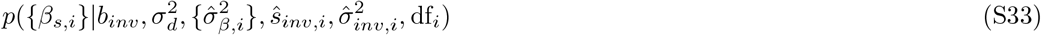

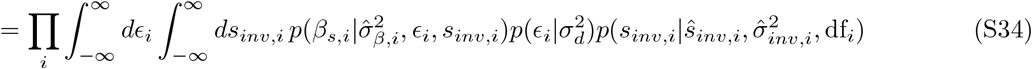

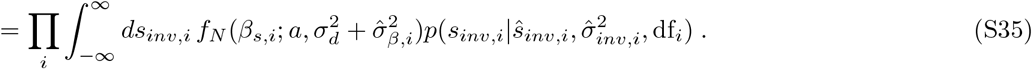

We numerically integrate the final step; we discretize *s*_*inv,i*_ into a grid of 10^5^ points, distributed equally between 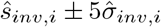.

##### Model 3

Similar to model 2, we now include *s*_*high,i*_ as a covariate,

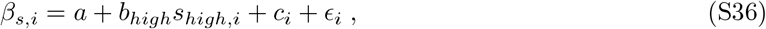

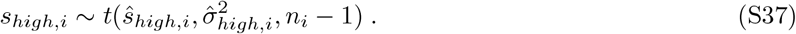

The likelihood of model 3 is analogous to that of model 2.

###### Evaluating model performance

We obtained parameter estimates for all three models by maximizing their respective likelihoods using `scipy.optimize.minimize`. We partition the variance in *β*_*s,i*_ across measurements, var(*β*_*s,i*_), into a maximum of three components: variance explained by the invasion or high-frequency fitness effects 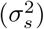, measurement error 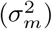, and unexplained variance 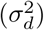. The estimate 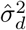 is directly obtained from the maximum likelihood estimate. We estimate 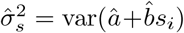, for each the invasion or high-frequency fitness effects. Then the variance associated with total measurement error is calculated as the remaining variance, 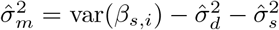. Results are summarized in Table S2.

**Table S2:** Results from ANCOVA models. Parameters fit via maximum likelihood. We find that Model 2 is the best fit to our data, both from its low Aikake Information Criteria (AIC), and from pairwise likelihood ratio tests (*p <* 10^−4^).

| Model | $\hat{a}$ | $\hat{b}$ | $\hat{\sigma}_d^2$ | $\hat{\sigma}_m^2$ | $\hat{\sigma}_s^2$ | $\log \hat{L}$ | AIC |
| --- | --- | --- | --- | --- | --- | --- | --- |
| 1 | -0.12 |  | 0.012 | 0.0076 |  | 35.8 | -67.6 |
| 2 | -0.029 | -0.26 | 0.0062 | 0.008 | 0.00554 | 43.4 | -80.9 |
| 3 | -0.11 | -0.057 | 0.012 | 0.0061 | 0.0021 | 35.2 | -64.4 |

### S3 Within-cycle time-courses

#### S3.1 Coculture experiments

Previous competition experiments focused on solely measuring two time points, both at the end of a growth cycle, separated by one growth cycle. We reasoned that measuring the within-cycle dynamics may help us understand the frequency-dependent ecological dynamics of strain competition.

After the initial growth cycles of fluorescently tagged strains (as previously described in section S1.1), we combined the strains into two cocultures, setting the volumetric fraction of one strain to either 10% or 90%. These cocultures were grown for one additional 24-hour cycle in DM25 medium, after which we performed a flow cytometry measurement—this served as the baseline, or time 0. Immediately afterward, we prepared replicate cultures by diluting the overnight culture 1:100 into fresh DM25, vortexing thoroughly, and distributing 1mL aliquots into individual wells of a glass 96-well plate. Each condition was performed in triplicate (three biological replicates). The plate was incubated in a 37°C warm room, shaking at 180rpm. At regular intervals (approximately every 45 minutes) over a 10-hour period, we briefly removed the plate to collect 60µL subsamples for flow cytometry analysis, capturing the exponential phase and the onset of stationary phase. An additional sample was collected at 24 hours to mark the end of the growth cycle. Subsamples were discarded after measurement, and all sampling times were recorded.

To understand the time-dependent growth dynamics, we sought to compute both the “instantaneous” growth rates and fitness effects of all competitions. We defined the growth rate (in units of per hour) at time *t* as simply log(*n*_*t*+Δ*t*_) − log(*n*_*t*_), where Δ*t* is the interval of time between measurements, and *n*_*t*_ is the total number of observed cells, after accounting for the measurement dilution factor. Analogously, we defined the fitness effect as logit(*f*_*t*+Δ*t*_) − logit(*f*_*t*_), where *f*_*t*_ is the frequency of the mutant relative to REL606. As computing derivatives is often quite noisy, we averaged over *n*_*t*_ and *f*_*t*_ for each time point *t* for all three biological replicates per condition. We computed the time-dependent growth rates and fitness effects via `np.gradient`, and then applied a simple convolution/moving average to further reduce noise associated with computing derivatives–for an estimate *x*_*t*_, we give the updated estimate as 0.5*x*_*t*_ + 0.25*x*_*t*−1_ + 0.25*x*_*t*+1_; the left boundary is given by 0.75*x*_*t*_ + 0.25*x*_*t*+1_, with an analogous equation for the right boundary. We then used standard bootstrapping–resampling with replacement–to compute the sampling errors associated with our estimates of growth rates and fitness effects.

#### S3.2 Measuring growth curves via plate reader

We sought to measure growth curves of all strain in monoculture. We prepared cultures as previously described, with six biological replicates per strain. To measure growth curves, we prepared 100µL cultures in DM25 in a flat bottom 96 well plate, and measured OD600 absorbance in a shaking plate reader (SpectraMax 190; Molecular Devices) over the course of 20 hours. We extracted an estimate of the average growth rate, *r*, by fitting an exponential curve to the portion of the growth curve in exponential growth, via standard ordinary least squares. We obtained an estimate by fitting the following model to the portions of the growth curves between the end of exponential phase and the beginning of stationary phase (5-10 hours),

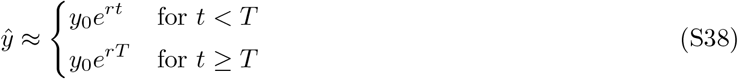

Here *y* is OD600 and *t* is time. We fit the parameters *y*_0_, *r*, and *T* simultaneously using ordinary least squares. Results are shown in Figure S20.

### S4 Batch culture resource competition model

As stated in the main text, we consider a simple model that tracks the dynamics of strain abundance, *n*(*t*), and resource abundance, *R*(*t*),

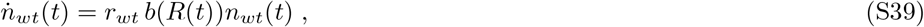

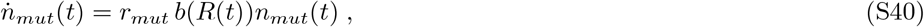

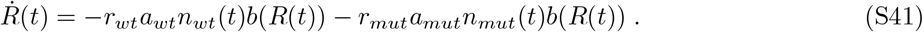

We will first choose *b*(*R*(*t*)) to be a simple step function *θ*(*R*(*t*)) for our analytical calculations. We also simulate the dynamics under other choices of *b*(*R*(*t*)) (Figure S17-S18). We are able to analyze the dynamics analytically in two limits, when *n*_*wt*_(*t*) ≫ *n*_*mut*_(*t*) and *n*_*wt*_(*t*) ≪ *n*_*mut*_(*t*). We call whichever strain that is in the majority, *maj*; in the limit that *f*_*maj*_ → 1, the resource dynamics reduce to,

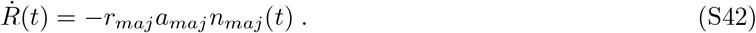

Integrating yields,

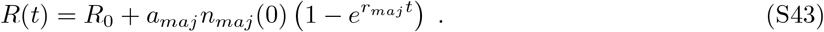

We can then find *T* where *R*(*T* ) = 0,

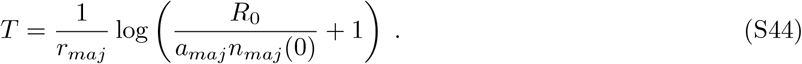

Fitness effects are similarly easy to calculate,

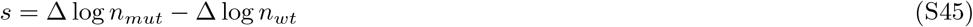

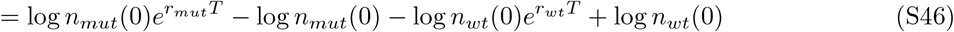

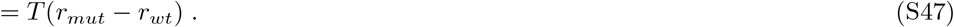

We can in fact obtain exact solutions to the ODEs for many monotonically increasing *b*(*R*), again in the limits *n*_*wt*_(*t*) ≫ *n*_*mut*_(*t*) or *n*_*wt*_(*t*) ≪ *n*_*mut*_(*t*). To see that, we note that in those limits,

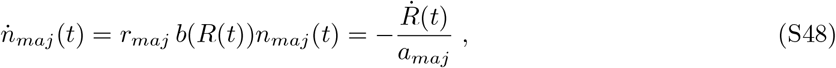

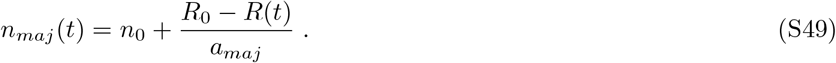

So, the system can be reduced to a single ODE,

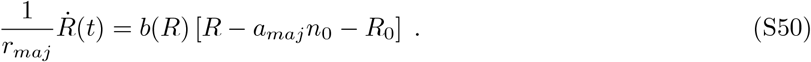

This equation can be exactly solved for *b*(*R*) = *R*^*n*^ for all positive integers *n*, for Monod’s equation 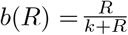, and various other choices (e.g. 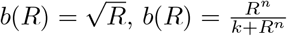, etc). This equation makes it clear that the timescale of resource depletion will always depend inversely on the growth rate of the strain in the majority, 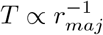, because we will always be able to rescale time by *r*_*maj*_; this is consistent with experimental plate reader measurements (Figure S20).

In the case that *b*(*R*) = *R* (the choice of MacArthur’s consumer-resource model), the population abundance dynamics follow a logistic equation,

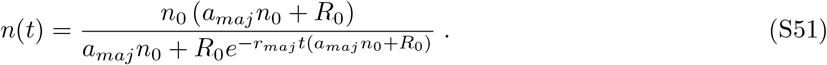

#### S4.1 Non-transitivity

Our model predicts that fitness effects will generally not be transitive across environments. We can calculate the invasion and high-frequency non-transitivities under our resource competition model (equations S7-S10),

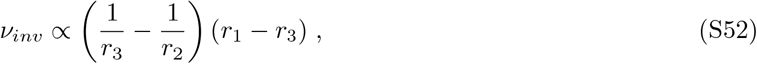

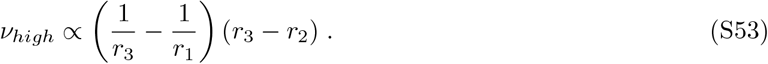

It is straightforward to calculate the expected non-transitivity under the other definitions of non-transitivity (equations S11-S18). As we see, the magnitude of non-transitivity generally increases as the difference between specific pairs of growth rates increases. Invasion and high-frequency non-transitivity will generally be non-zero unless either *r*_3_ = *r*_2_ or *r*_1_ = *r*_3_.

#### S4.2 Frequency-dependent slope

We can compute the expected relationship between the invasion fitness of a mutant, *s*_*inv*_, and the frequency-dependent slope (Figure S19). Here, we simply take the frequency-dependent slope to be Δ*s* = *s*_*high*_ − *s*_*inv*_. This differs from the frequency-dependent slope calculated from our data, *β*_*s*_–while *β*_*s*_ was computed as the slope of a fitted linear regression to the data, Δ*s* is computed by taking the slope between the fitness effects as the frequency approaches zero or one. If the form of *s*(*f* ) is actually linear, then the two slopes should be equivalent to each other, *β*_*s*_ = Δ*s*. Deviations from linearity will, in general, cause *β*_*s*_ ≠ difference in *β*_*s*_ and Δ*s* will presumably be small, assuming that *s*(*f* ) is not ill-behaved.

For generality, we lump the prefactor of *T* into a new constant, *c*, so that *T* = *c/r*_*maj*_. Following some algebra, we calculate Δ*s* as, Δ*s*; however, the

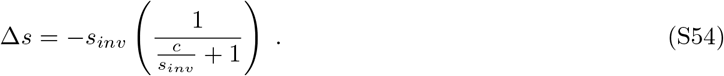

This expression can be further simplified in two limits. When *s*_*inv*_ is large and positive, *c* ≪ *s*_*inv*_, the slope will be linearly related to the invasion fitness,

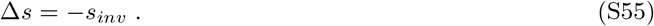

In the opposite limit, *c* ≫ *s*_*inv*_,

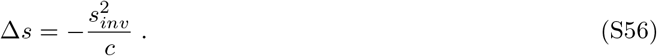

### S4.3 Comparing slopes of *s* **and** *w*

Under our model, *s* will generically be frequency-dependent, because of it’s dependence on the frequency-dependent time to depletion, *T* . However, *w* is a ratio between two unitless variables (equation 2), and is therefore also unitless; *w* will therefore be unaffected by the frequency-dependent time to depletion. For example, in our model under the simple case where *b*(*R*) = *θ*(*R*), *w* will be,

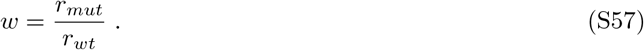

We can thus compare the frequency-dependence of *w* and *s* to determine what proportion of frequency-dependence is due to the resource depletion effect demonstrated in our model. Specifically, we consider the simple case where there are two possible sources of frequency dependence–resource depletion, and frequency-dependent growth rates. The growth rates of the mutant and wild-type may differ arbitrarily between the high- and low-frequency conditions; we call the growth rates 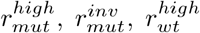, and 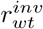. We write the low-frequency growth rates in terms of the high-frequency growth rates,

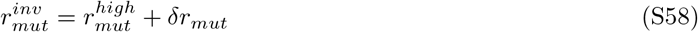

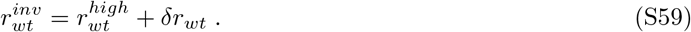

We make the simplifying assumption that we can approximate the frequency-dependent slope as the difference between the high- and low-frequency fitness effect. Again, for generality, we lump the prefactor of *T* into a new constant, *c*, so that *T* = *c/r*_*maj*_. Now, the predicted frequency-dependent slope of *s* is (dropping superscripts),

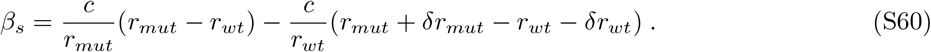

The corresponding frequency-dependent slope of *w* is,

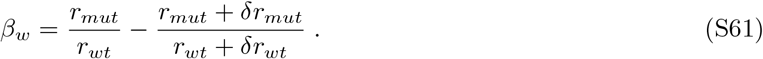

Following some algebra, we find a relationship between *β*_*s*_ and *β*_*w*_,

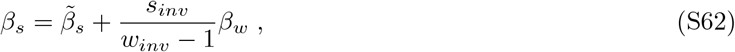

where 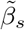 is the slope of *s* if resource depletion were the only source of frequency dependence, i.e. if *δr*_*mut*_ = *δr*_*wt*_ = 0. We compute 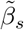 as a measure of how much resource depletion contributes to frequency-dependent slopes in our experimental data (Figure S16).

---

**Figure S1:**
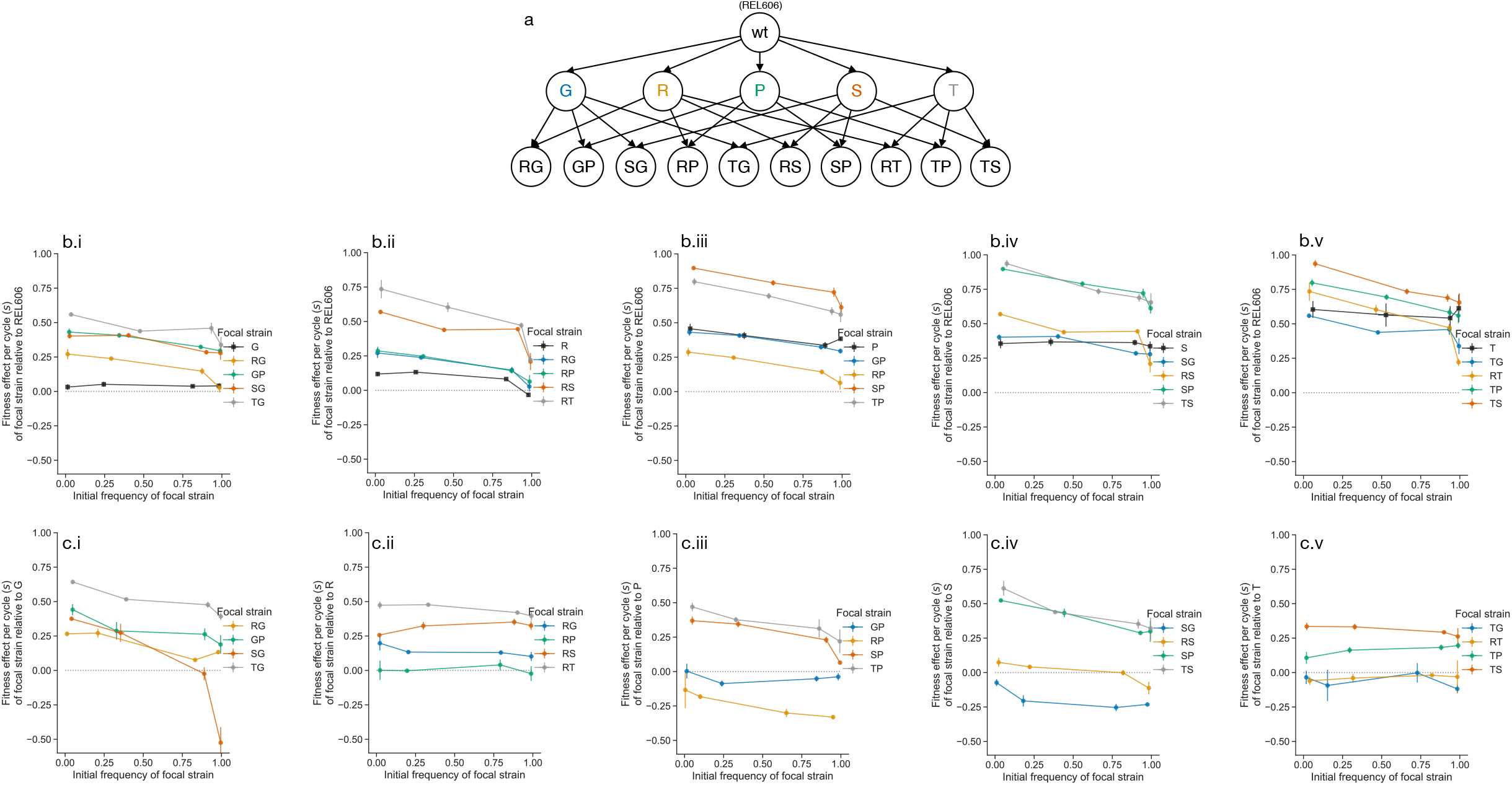
Ubiquity of frequency-dependence. Identical to Figure 1; linear x-axis instead of a logit x-axis. (**a**) We study a set of single and double mutants, all derived from the ancestor of the *E. coli* LTEE, REL606. (**b**) We measured the fitness effects, *s*, of all single and double mutants against REL606 as a function of frequency. Each subpanel (i-v) shows the measured frequency-dependent fitness effects for each single mutant and derived double mutants. Note that we plot the frequency-dependent fitness effects of double mutants twice–once on each subpanel corresponding to an immediate ancestral single mutant–to allow for comparison. (**c**) We additionally measured the frequency-dependent fitness effects of all double mutants against their single mutant ancestors. Error bars represent standard errors across biological replicates (*n* =4∼6).

**Figure S2:**
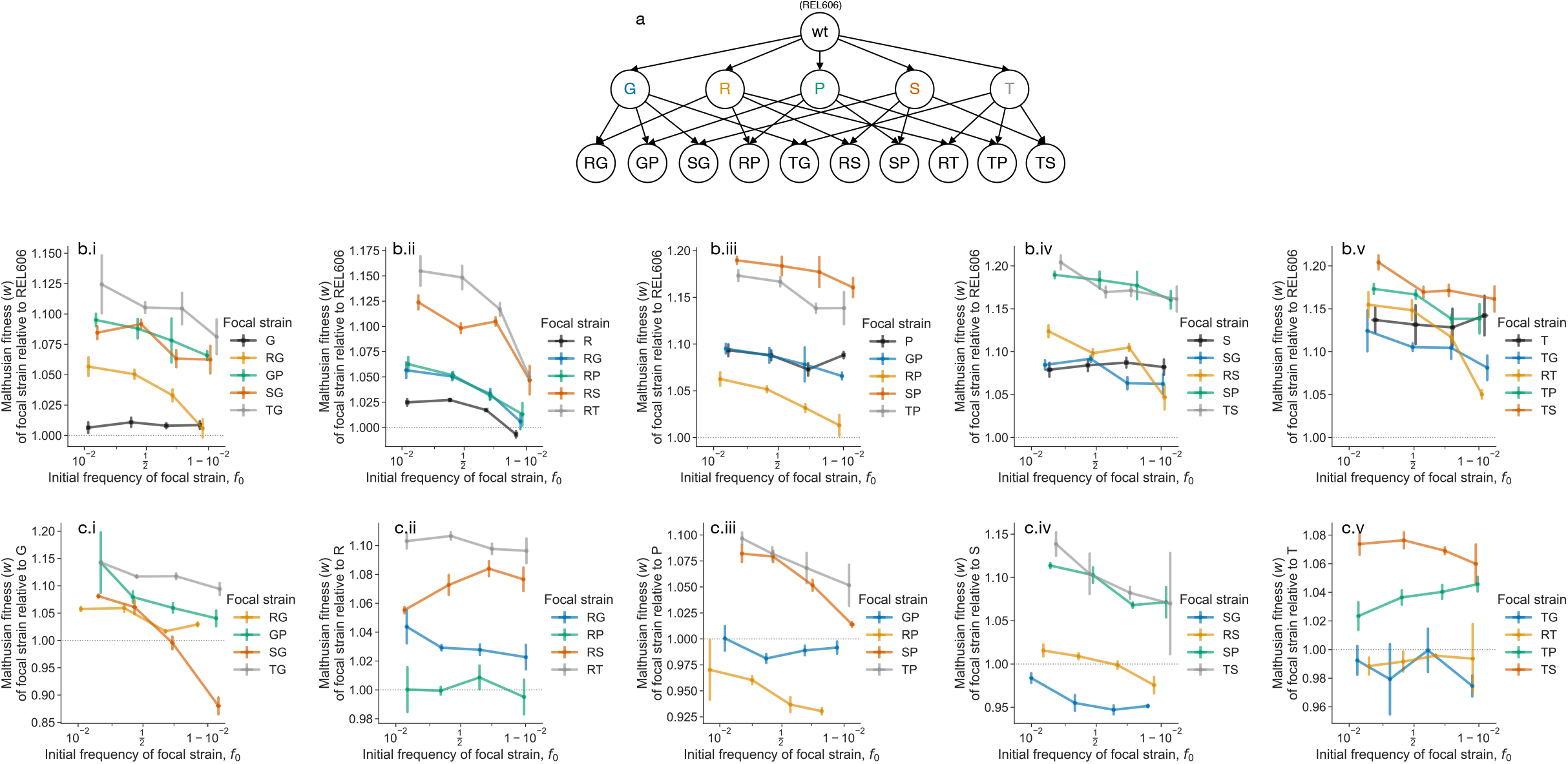
Frequency-dependence with Malthusian fitness. Similar to Figure 1, except fitness is calculated as the ratio of Malthusian parameters (*w*) instead of selection coefficient (*s*). (**a**) We study a set of single and double mutants, all derived from the ancestor of the *E. coli* LTEE, REL606. (**b**) We measured the fitness effects, *s*, of all single and double mutants against REL606 as a function of frequency. Each subpanel (i-v) shows the measured frequency-dependent fitness effects for each single mutant and derived double mutants. Note that we plot the frequency-dependent fitness effects of double mutants twice–once on each subpanel corresponding to an immediate ancestral single mutant–to allow for comparison. (**c**) We additionally measured the frequency-dependent fitness effects of all double mutants against their single mutant ancestors. Error bars represent standard errors across biological replicates (*n* =4∼6).

**Figure S3:**
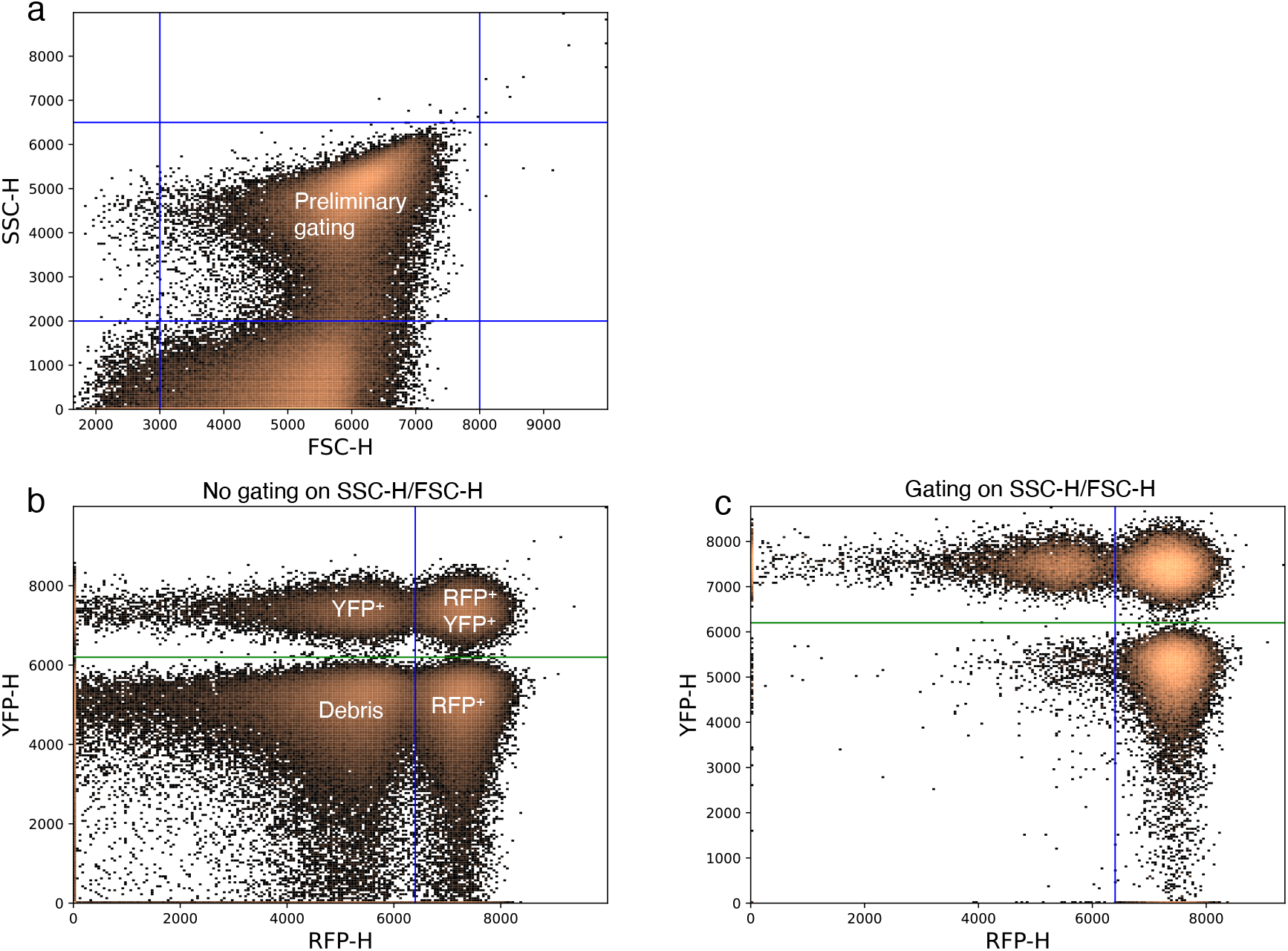
Example of flow cytometry gating strategy. (a) We first gate on FSC-H/SSC-H to select for the cells present in our sample, and deplete debris. (b) In the absence of the initial FSC-H/SSC-H, we see a large cloud of debris particles in the bottom left corner, which (c) largely disappears after the initial gating. We use threshold gates to divide the population into YFP^+^, RFP^+^, and double positive events. The double positive events correspond to coincident events (which are difficult to eliminate due to the small size of *E. coli* ). We use a previously validated pipeline [1] to estimate the frequency of each fluorescently labeled subpopulation, accounting for coincident events.

**Figure S4:**
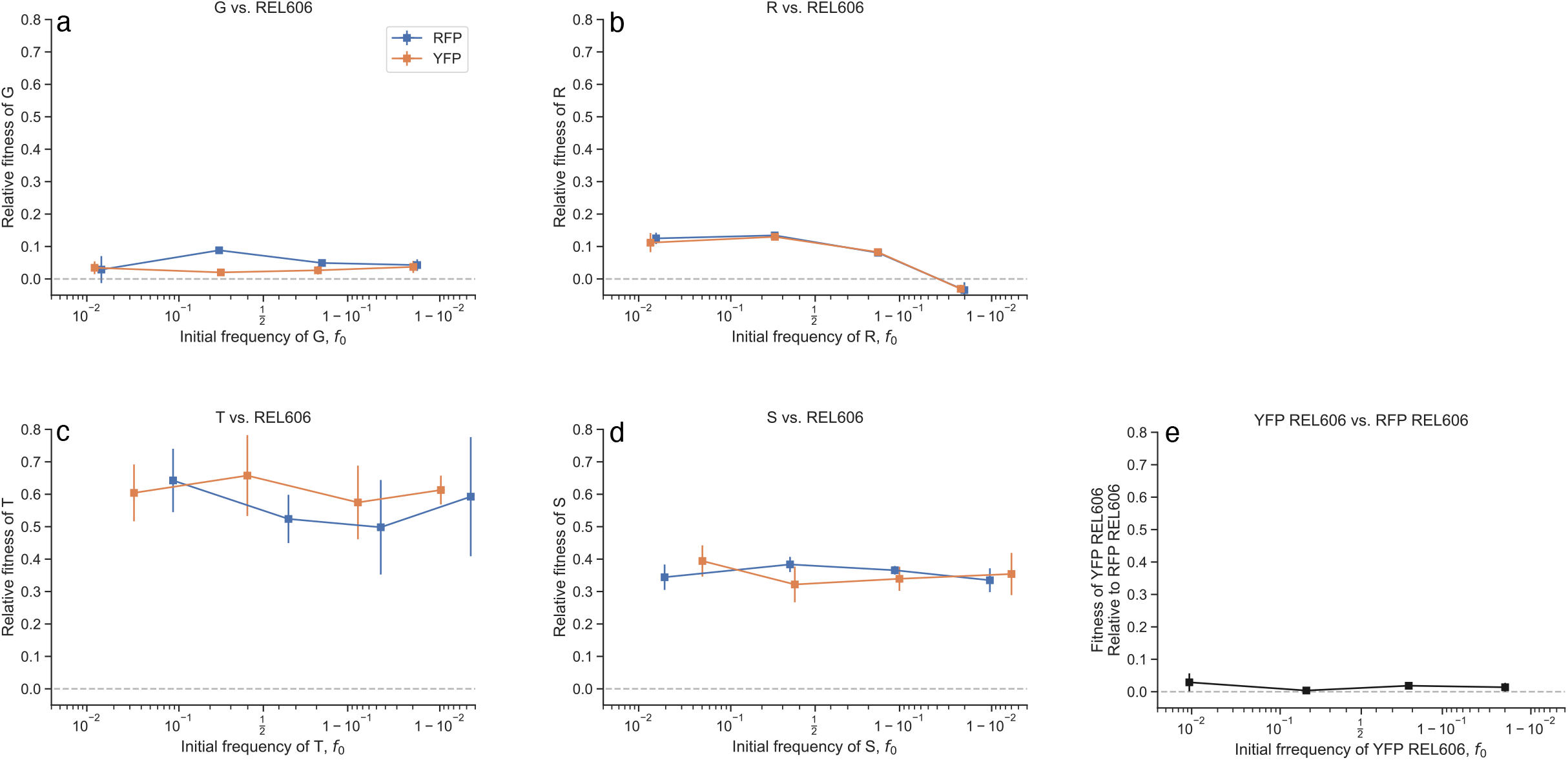
Fluorophores do not differentially impact fitness. (a-d) We conducted several control experiments where we competed a either an RFP-tagged mutant against a YFP-tagged REL606 clone, or vice-versa. We measured the fitness effects, *s*, of all mutants against REL606 as a function of frequency. Frequency-dependent fitness effects after switching fluorophores are generally within error bars. (e) Similarly, we competed a YFP-tagged REL606 clone against an RFP-tagged REL606 clone; fitness effects across all measured frequencies are not significantly different from zero. Error bars represent standard errors.

**Figure S5:**
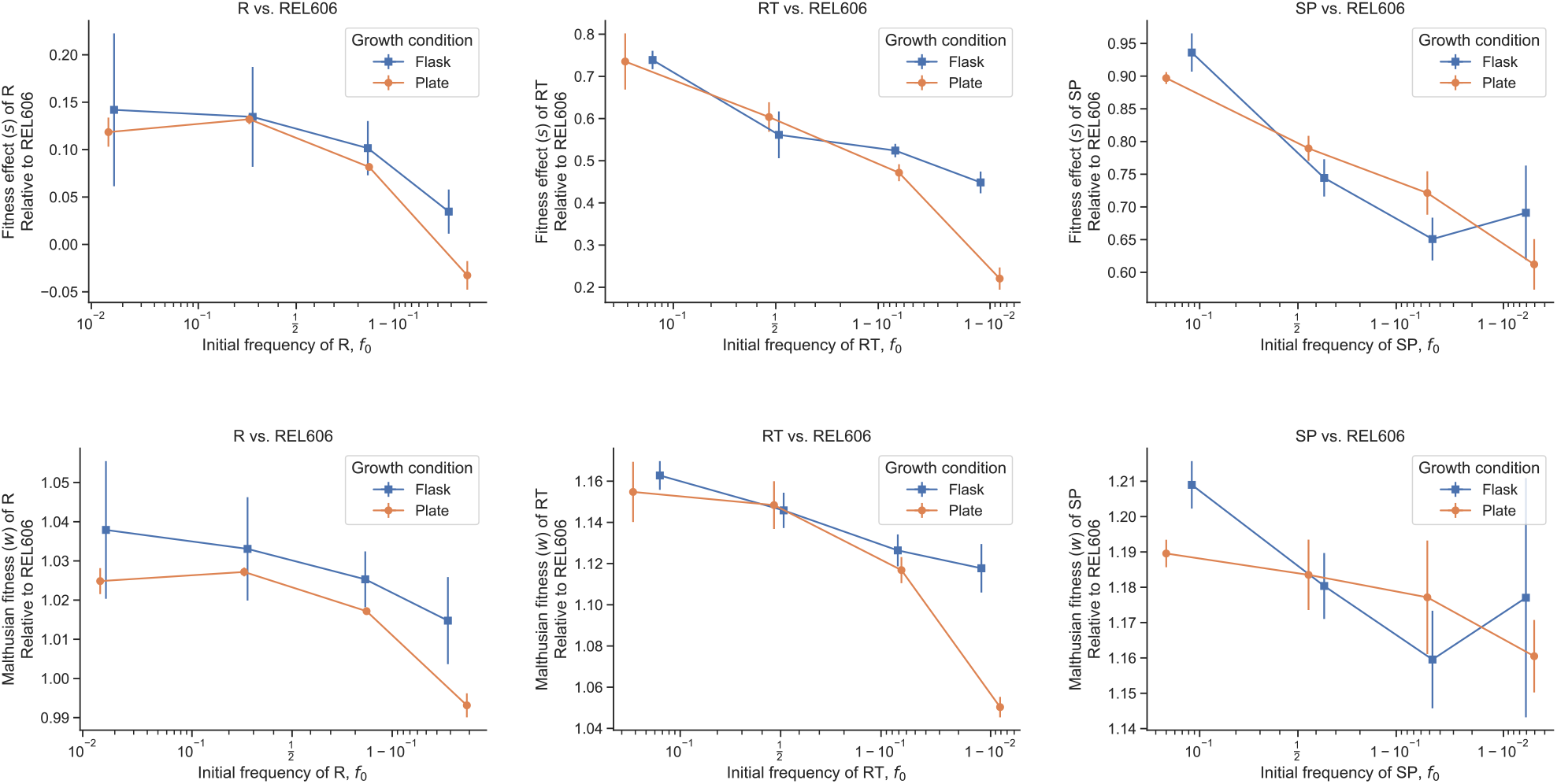
Comparing fitness effects in flasks and glass 96 well plates. In order to investigate if there are any significant differences between fitness effects in glass 96-well deep well plates (where we did the majority of our fitness measurements) and glass flasks (the home environment of the LTEE), we repeated several competition experiments in glass flasks. We computed fitness as both the slope of logit frequency (*s*; top row) and the ratio of Malthusian parameters (*w*, bottom row). The *w* fitness measurements appear noisier than *s*, primarily because *w* uses population size measurements, which are generally noisier than frequency measurements in the flow cytometer. For all three competitions, we generally get quantitative agreement between the measurements in flasks and plates. Overall, this data supports the claim that the glass plates generally recapitulate the environment in the flask.

**Figure S6:**
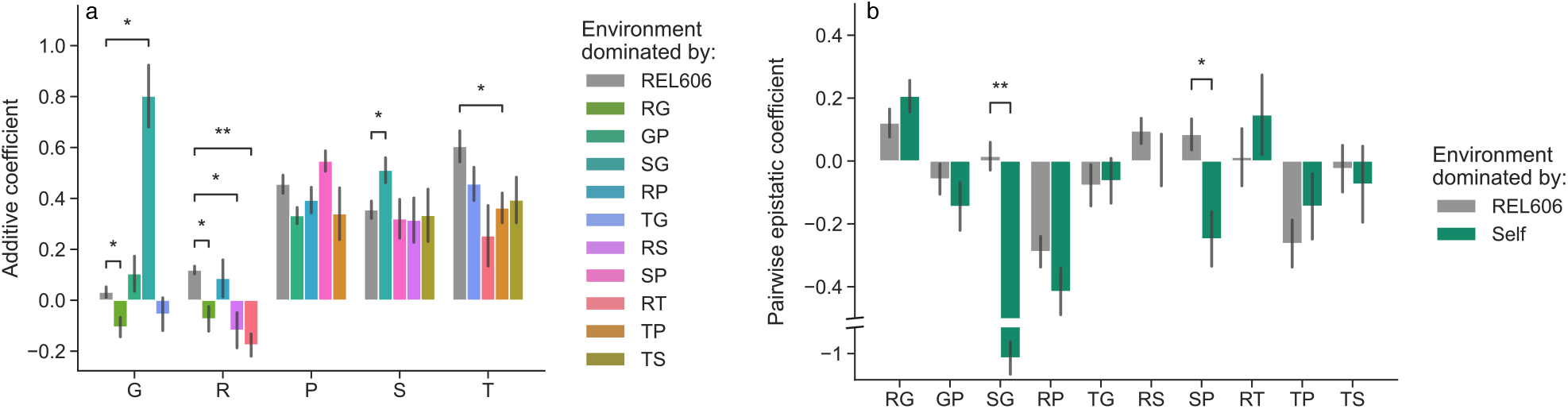
Frequency-dependent epistasis. Estimation of (a) additive and (b) epistatic coefficients of double and single mutants. Coefficients are calculated using fitness effects at either invasion or high frequencies; we compare coefficients when different clonal strains are in the vast majority of the population, and thus dominating the environment. Coefficients are calculated using the methods presented in section S2.3. * *p <* 0.05, ** *p <* 0.01, *** *p <* 0.001, post-FDR correction.

**Figure S7:**
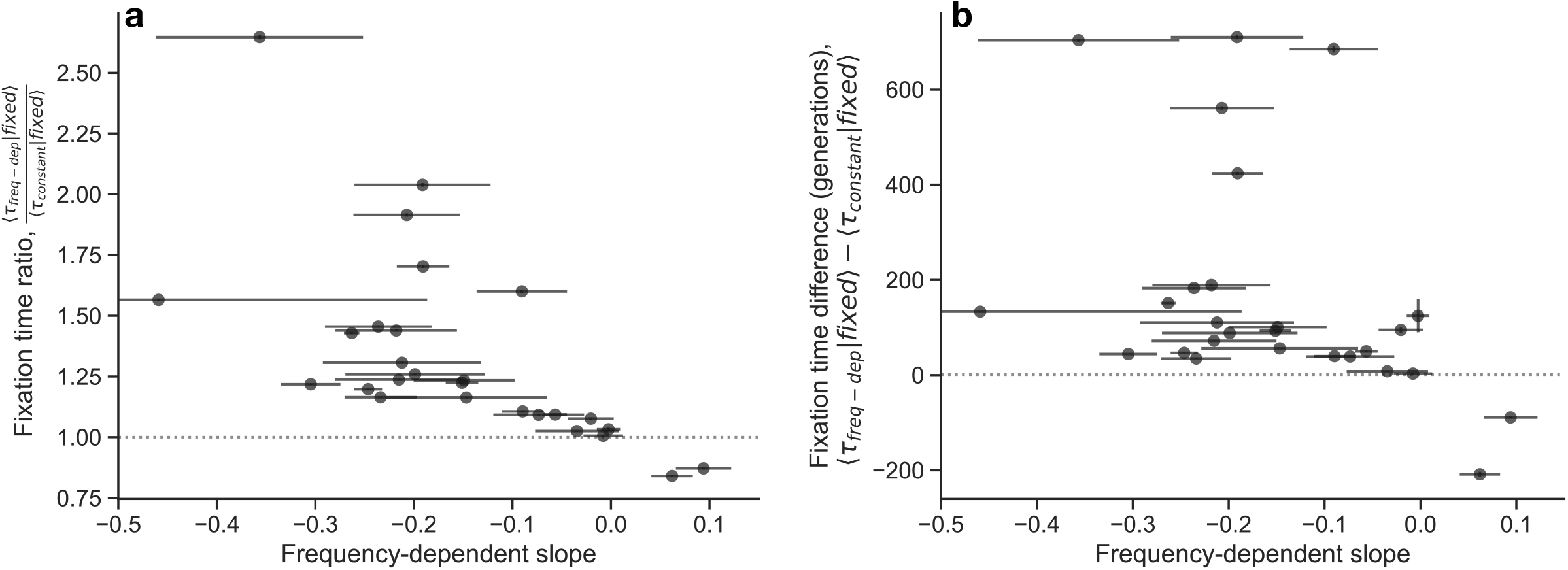
Comparison of simulated mutant fixation time estimates with and without frequency-dependent fitness effects. We numerically simulated the frequency (*f* ) dynamics of every strain pair in our dataset by numerically integrating the one-locus Langevin equation, 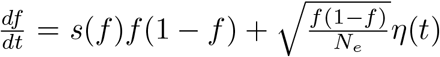, where *η*(*t*) is standard Gaussian white noise, and we set *N*_*e*_ = 10^6^. We either approximate *s*(*f* ) as a linear function with previously estimated parameters (see Figure 2 in the main text), or we set *s*(*f* ) to be constant (at the invasion fitness of the linear approximation). We simulate the frequency dynamics for each parameter set (corresponding to each strain competition) 2000 times, with the focal strain set to an initial frequency of 10^−6^; if the focal strain fixed, we recorded the time to fixation. We compare the time to fixation with frequency-dependent or constant fitness effects by either taking (a) the ratio of average fixation times or (b) the difference in average fixation times.

**Figure S8:**
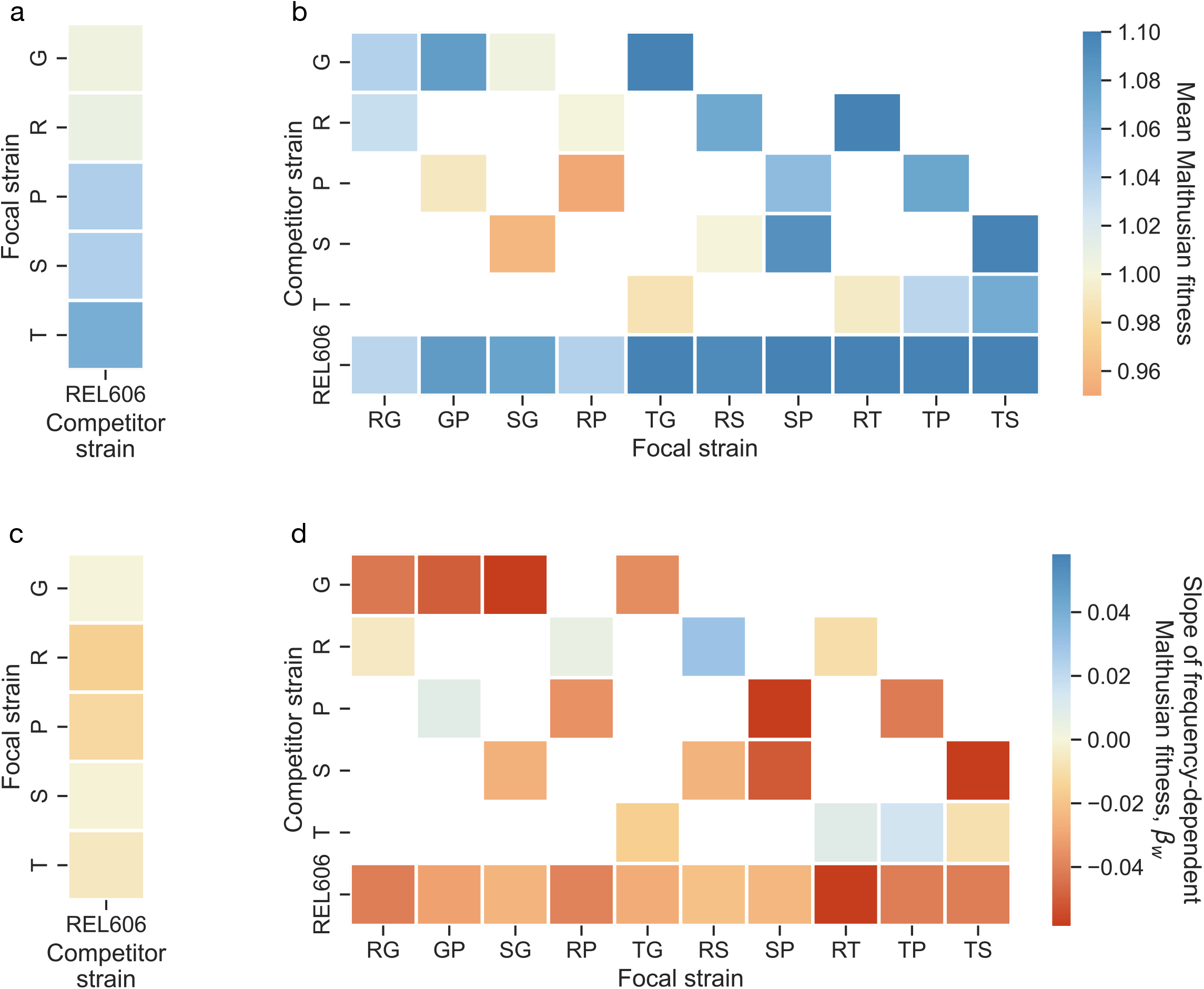
Summaries of Malthusian fitness. Identical to Figure 2, except using Malthusian fitness *w* instead of *s*. (**a-b**) Mean fitness effect of the focal strain relative to the competitor strain, across biological replicates and measured frequencies, of all competitions. (**c-d**) Mean slope of frequency-dependent fitness effects for all competitions, of the focal strain relative to the competitor strain. Slopes obtained through orthogonal distance regression (SI section S2.4).

**Figure S9:**
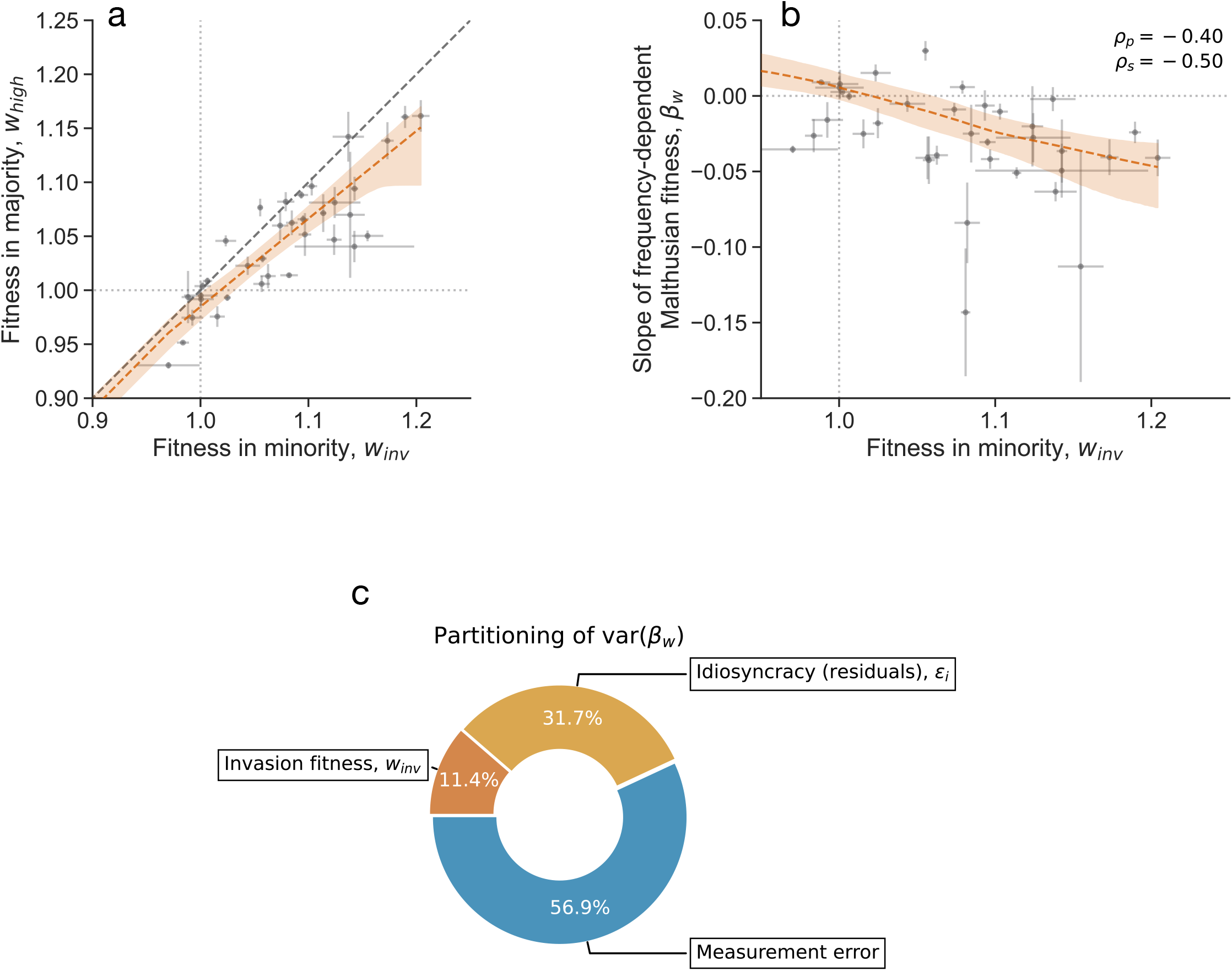
Statistical patterns of frequency-dependent fitness effects. Identical to Figure 3, except using Malthusian fitness *w* instead of *s*. (**a**) Comparison of low-frequency (*w*_*inv*_) with high-frequency (*w*_*high*_) Malthusian fitness for all competitions. (**b**) The average slope of frequency-dependent fitness effects is negatively correlated with the fitness effect of the focal strain in the minority. *ρ*_*p*_ represents Pearson’s correlation; *ρ*_*s*_ represents Spearman’s correlation. The red lines represent a LOWESS fit. All error bars represent standard errors. (**c**) Partitioning the variance in frequency-dependent fitness effect slopes using ANCOVA into three components: invasion fitness, biological idiosyncrasy, and measurement error. The higher proportion of variance in *β*_*w*_ attributed to measurement error is likely due to the fact that the *w* fitness measurement is noisier than that for *s*.

**Figure S10:**
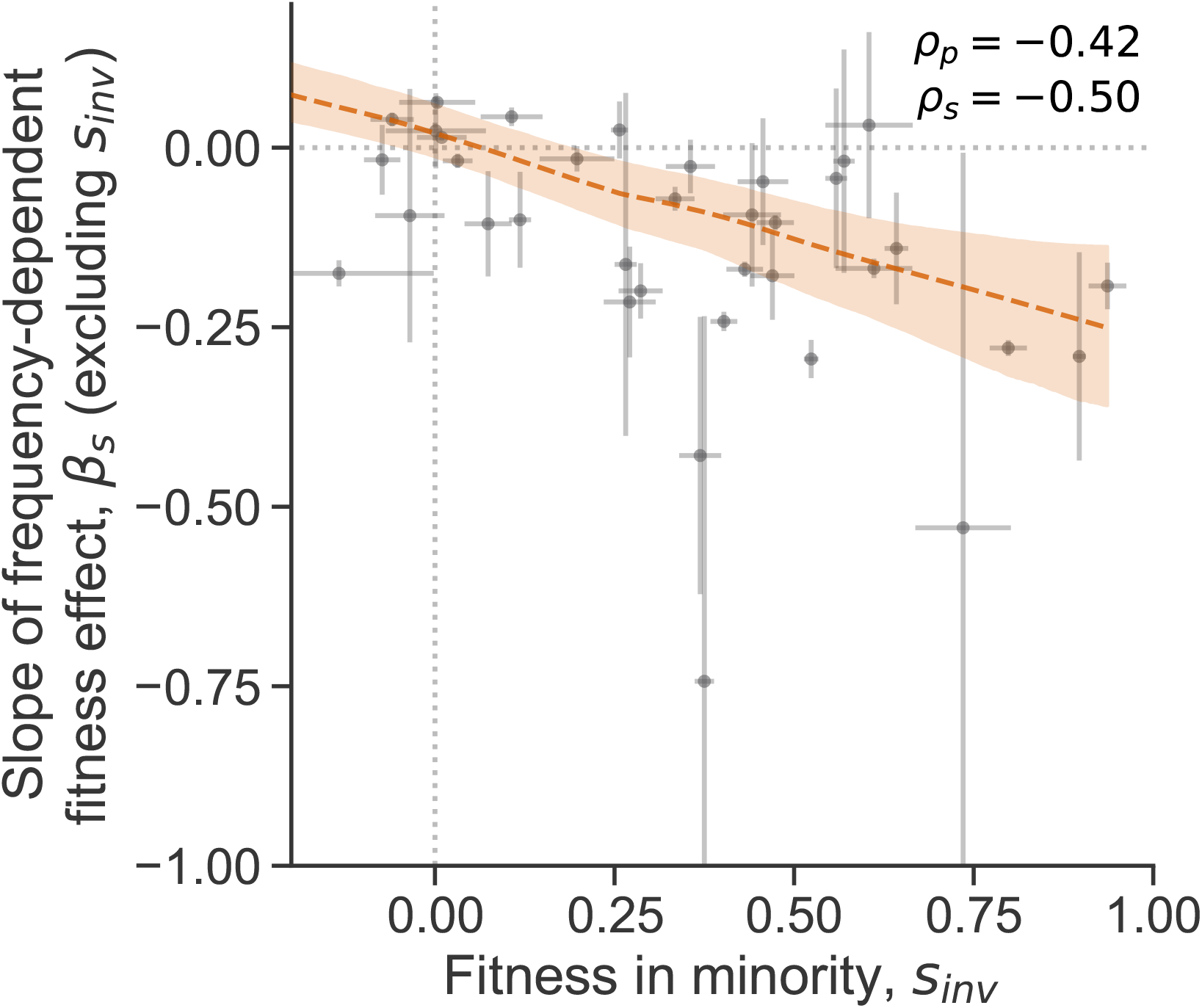
Frequency-dependent slopes. Identical to Figure 3a, except calculating the slope by using all points except those corresponding to the invasion fitness.

**Figure S11:**
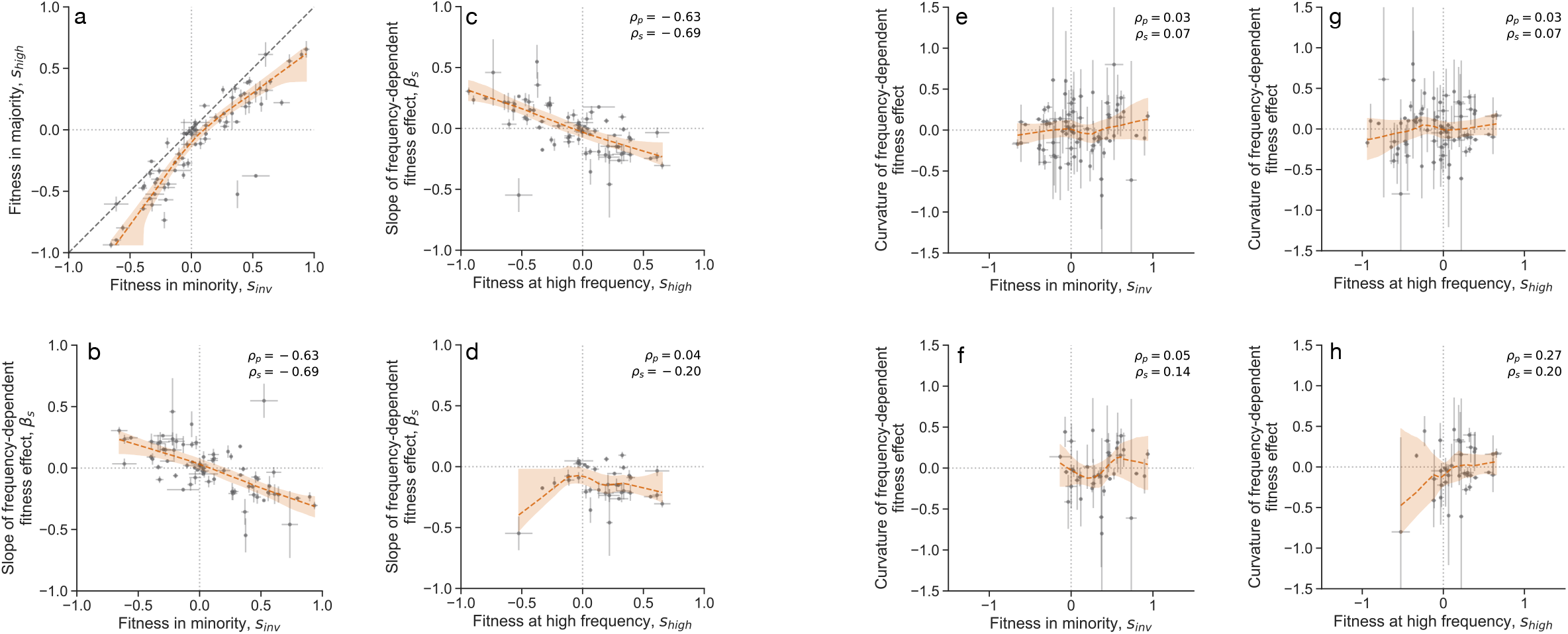
Statistical relationships between invasion/high-frequency fitness effects and frequency dependence (related to Figure 4). Specifically, predictors of (a-d) the average slope of the fitness effects, and (e-h) the curvature of fitness effects. (a-b) Analogous to Figure 4a-b, but we treat each experiment as two points, where either strain in the competition can be the focal or competitor strain. (c) Relationship between the fitness at high frequency and frequency-dependent slope; (d) analogous, but only considering points where the competitor strain is more ancestral relative to the focal strain. Relationship between curvature of frequency-dependence and (e-f) invasion fitness effect or (g-h) fitness effect at high frequency; either considering (e,g) points from both perspectives, or (f,h) only points where the competitor strain is more ancestral relative to the focal strain. Red lines represent LOESS regression fit; shaded region is a 95% confidence interval (obtained from bootstrapping). Error bars on points represent standard errors. Here, *ρ*_*p*_ is the estimated Pearson’s correlation coefficient; *ρ*_*s*_ is the estimated Spearman’s correlation coefficient.

**Figure S12:**
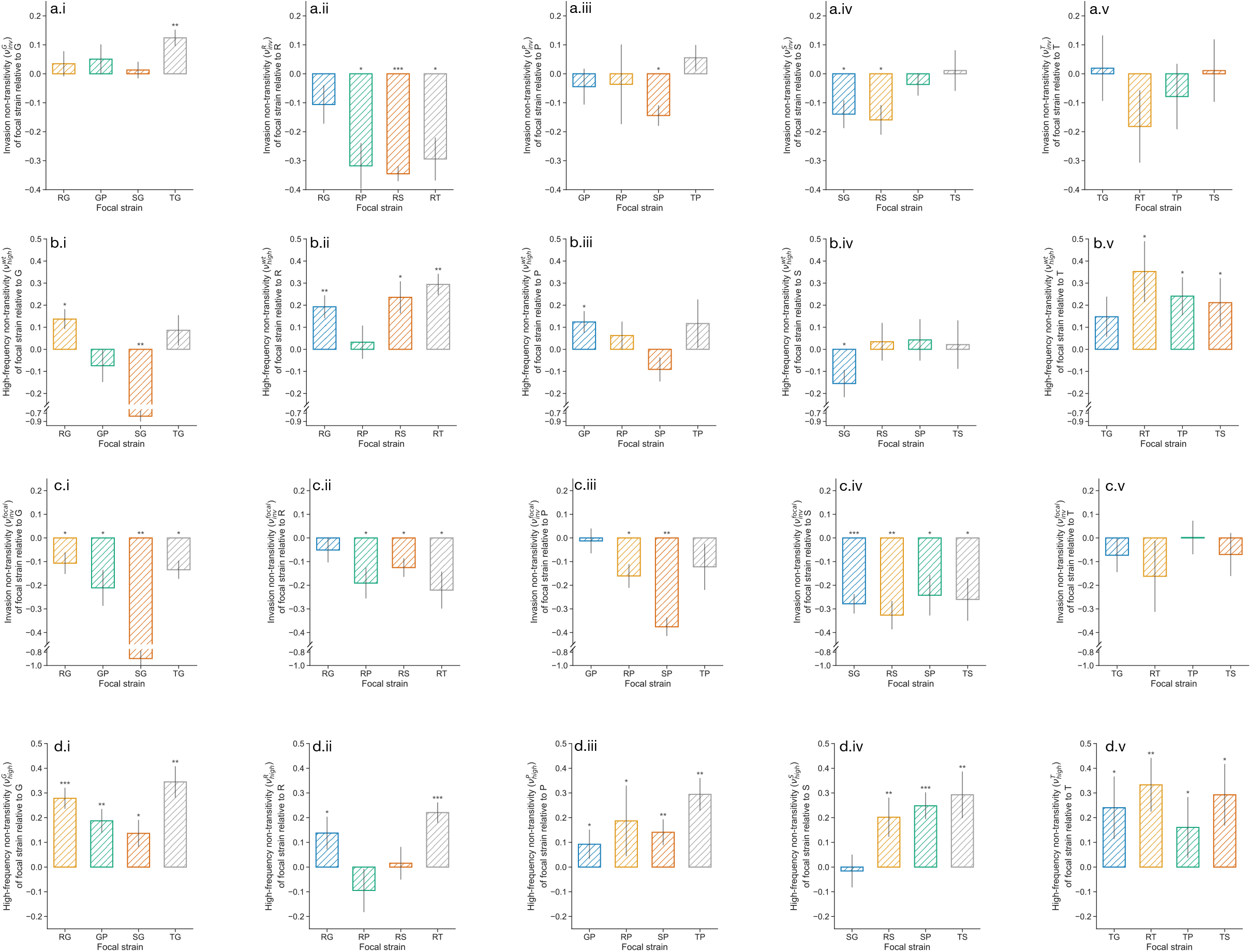
Non-transitivity of fitness effects (related to Figure 4). Quantification of non-transitivity using alternative definitions of frequency-dependent non-transitivity (equations S11-S18). Each row (a-d) represents a different definition of non-transitivity. Each column (i-v) represents non-transitivity of double mutants against different single mutants, (i) G, (ii) R, (iii) P, (iv) S, (v) T. Note that the y axis scales differ between rows. * *p <* 0.05, ** *p <* 0.01, *** *p <* 0.001, post-FDR correction.

**Figure S13:**
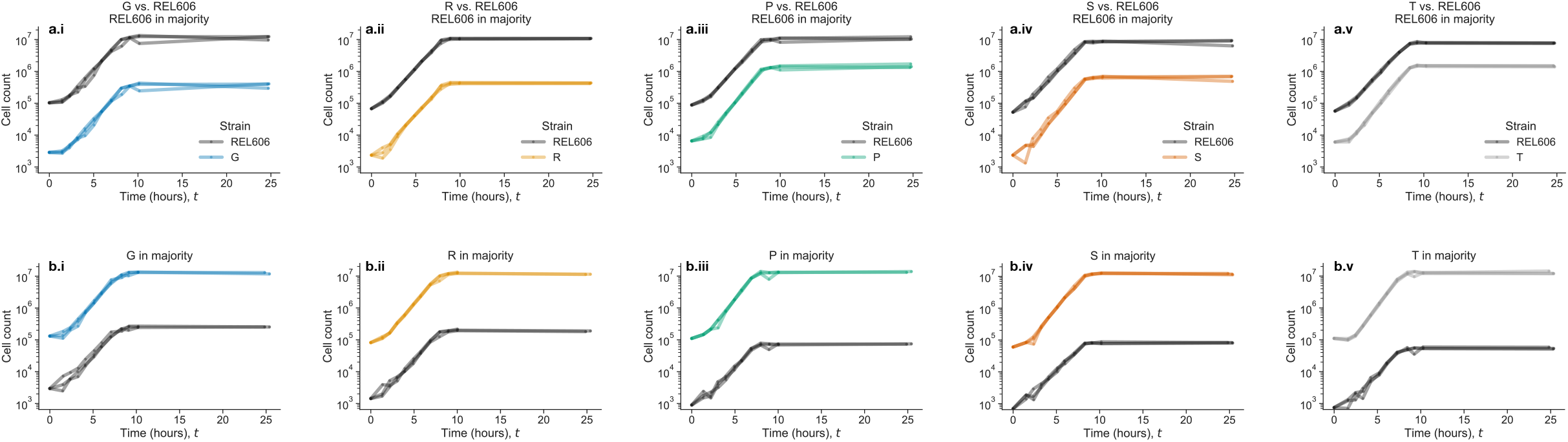
Within-cycle population dynamics, including the 24 hour time point (related to Figure 5). (a-b) Dynamics of cell counts over the course of the entire groowth cycle; where either (a) REL606 is in the majority of the population, or (b) the mutant is in the majority of the population. Trajectories are shown for all three biological replicates per condition. Population sizes are approximately constant from 10-24 hours, as expected.

**Figure S14:**
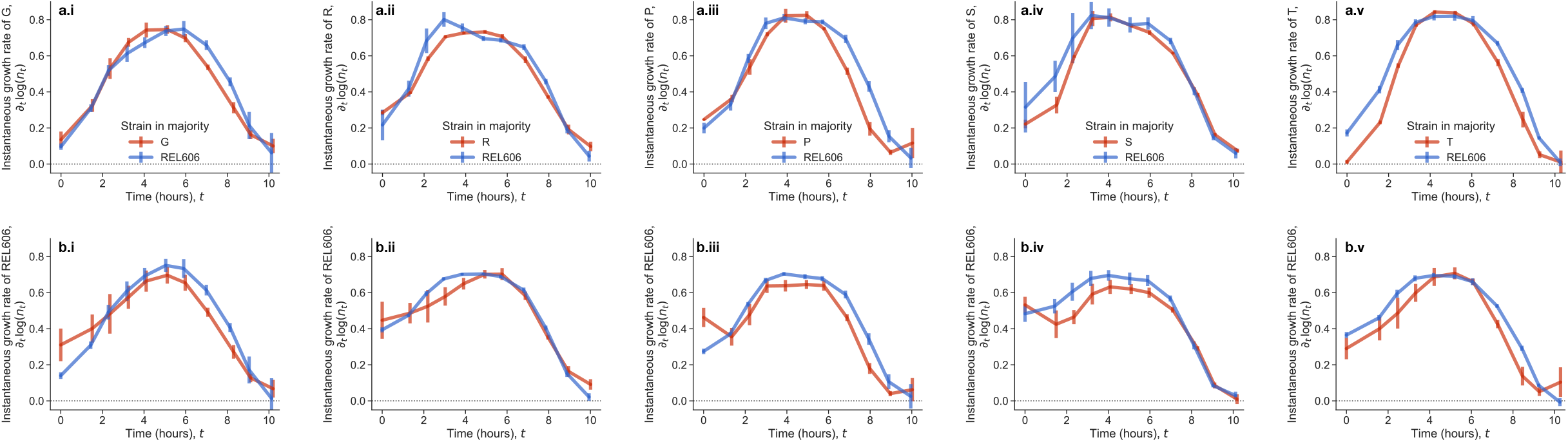
Within-cycle population growth rates (related to Figure 5). Estimates of the time-dependent growth rates of (a) the mutants and (b) REL606 in each experiment. Error bars represent standard errors.

**Figure S15:**
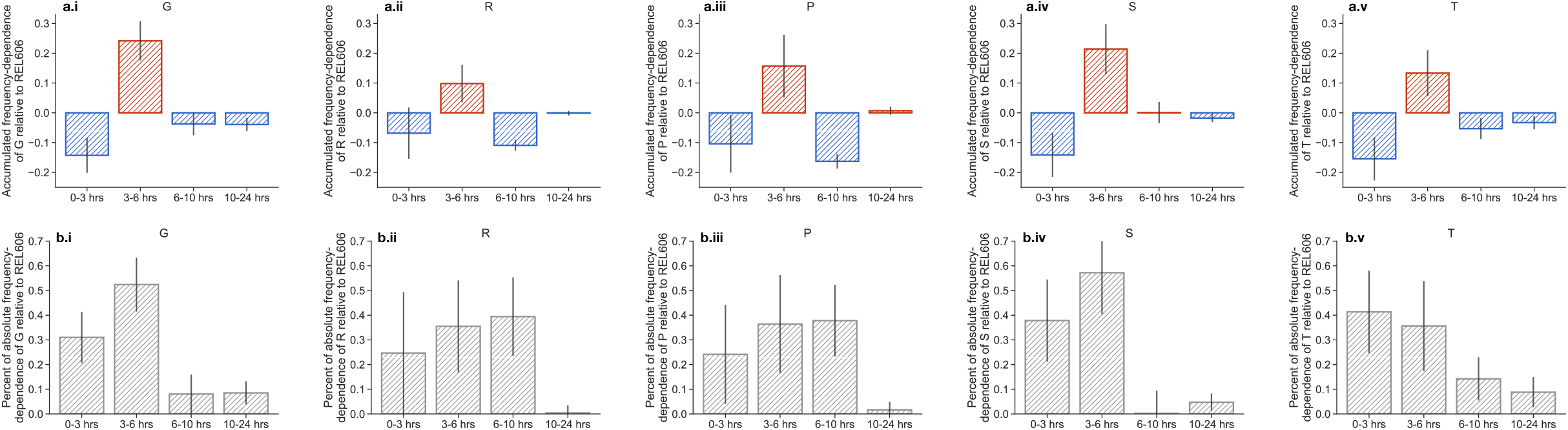
Accumulated frequency-dependence over time intervals (related to Figure 5d). (a) We subtracted the total accumulated fitness effects at each time interval to get an estimate of the time-dependence frequency-dependence, *s*_*high*_ − *s*_*inv*_ (fitness at high mutant frequency - low mutant frequency). (b) We computed the percent contribution of each time bin to the total absolute frequency-dependence, i.e. |*s*_*i,high*_ − *s*_*i,inv*_|∑_*j*_ |*s*_*j,high*_ − *s*_*j,inv*_|. Error bars represent standard errors.

**Figure S16:**
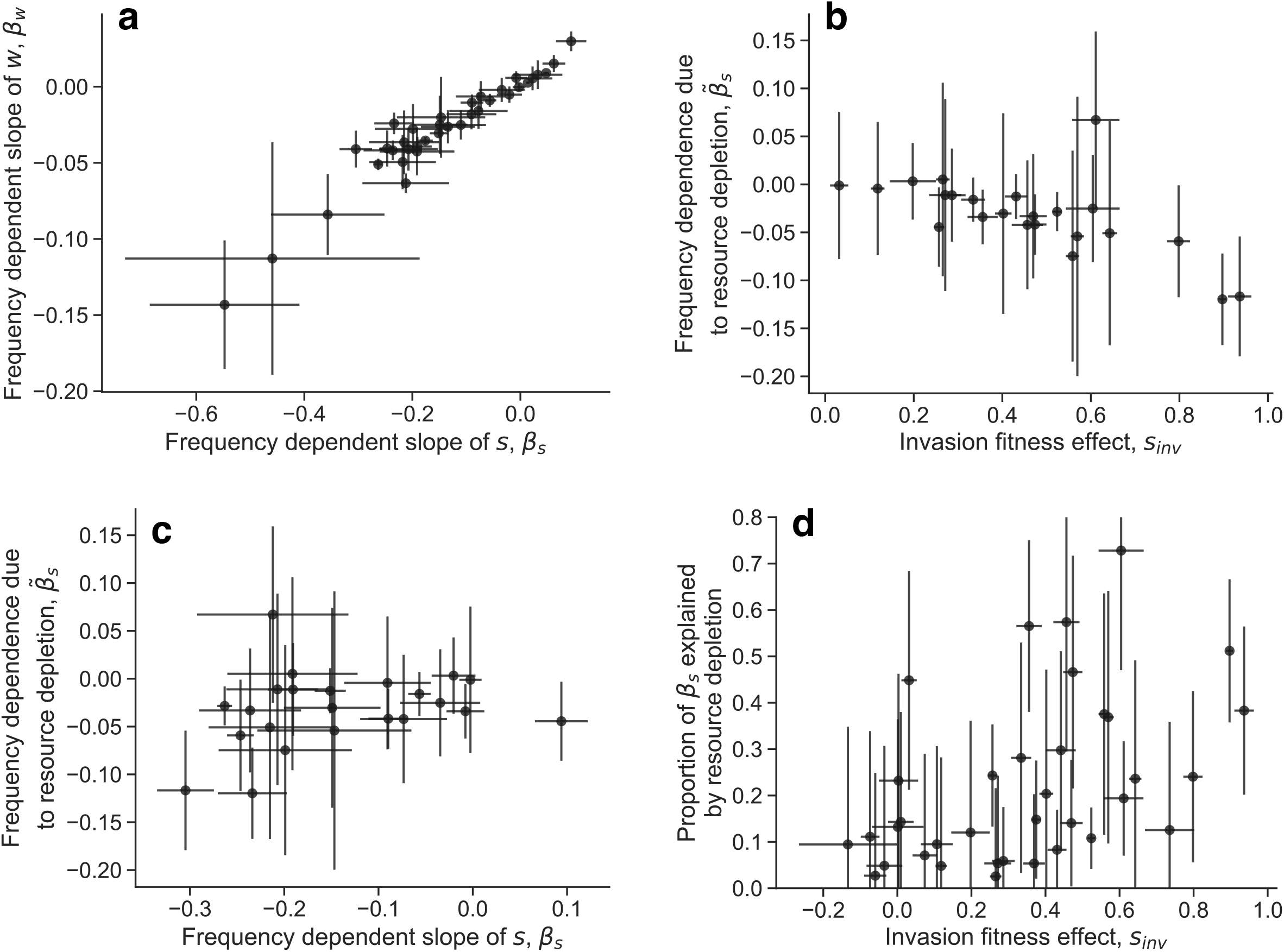
Comparing *s* and *w* slopes. (a) Relationship between *s* and *w* slopes across all competitions. (b-c) Comparison of the estimated contribution of resource competition to the frequency-dependent slope, 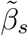 (equation S62), to (b) invasion fitness and (c) *β*_*s*_. (d) We compute the absolute proportion of *β*_*s*_ explained by resource competition, i.e 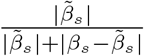. All error bars represent standard errors, obtained via standard bootstrapping. We excluded points in panels (b) and (c) with noisy estimates of 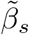, i.e. those with standard errors *>* 0.15.

**Figure S17:**
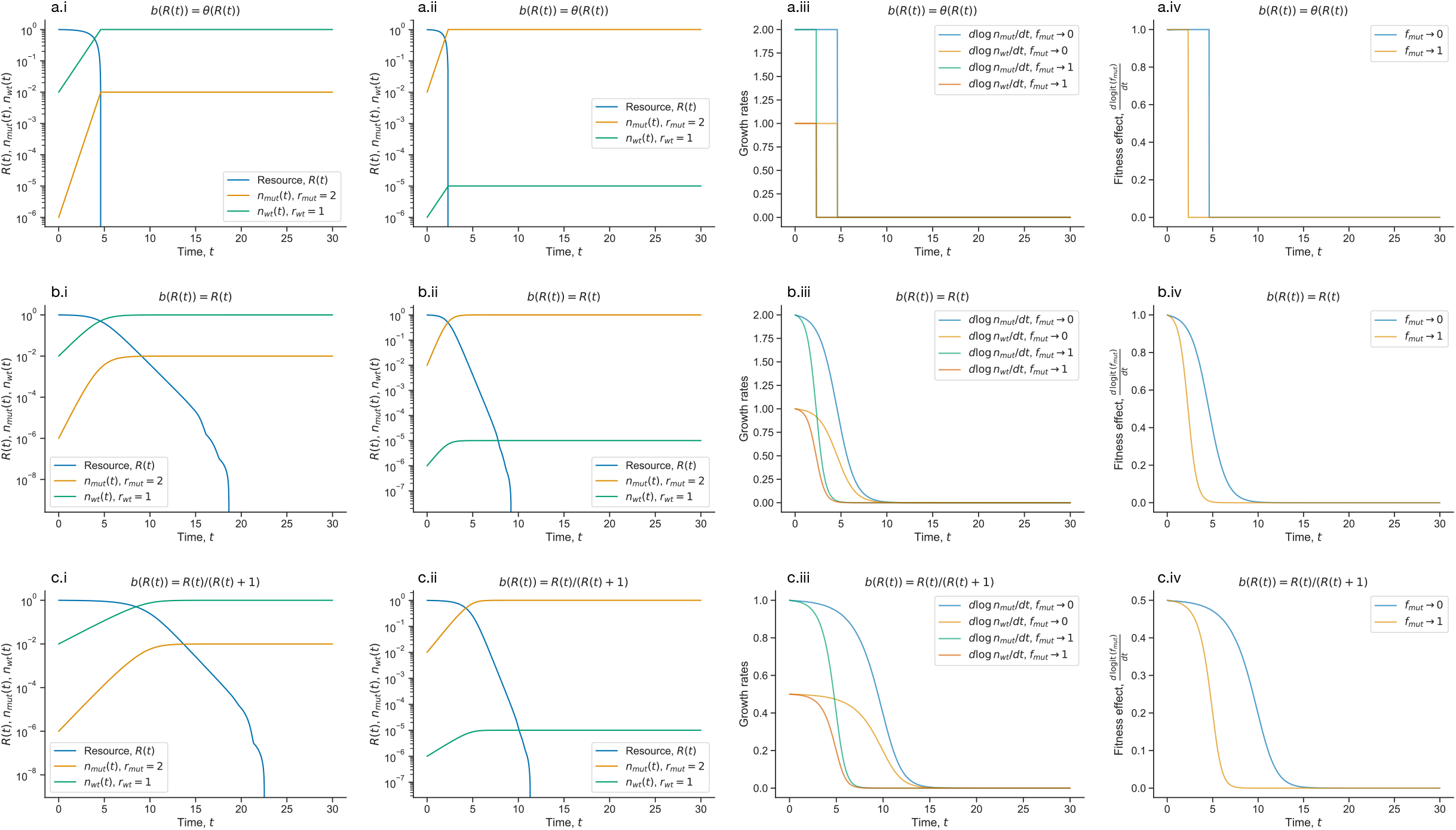
Examples of simulations of batch culture resource competition dynamics. Here we show examples of simulated batch culture resource competition dynamics, obtained by numerically solving equations 2-4 (in the main text). We choose three different values of *b*(*R*(*t*)), (a) a step function, (b) a linear function, and (c) a Monod function. We varied the initial frequency of the mutant strain, using either (i) *f* = 10^−4^ or (ii) *f* = 1 − 10^−4^. We also compute and compare (iii) population growth rates, along with (iv) instantaneous fitness effects. Parameter values: initial amount of resource, *R*_0_ = 1; initial total population size *N* = 10^−2^; *a*_*wt*_ = *a*_*mut*_ = 1.

**Figure S18:**
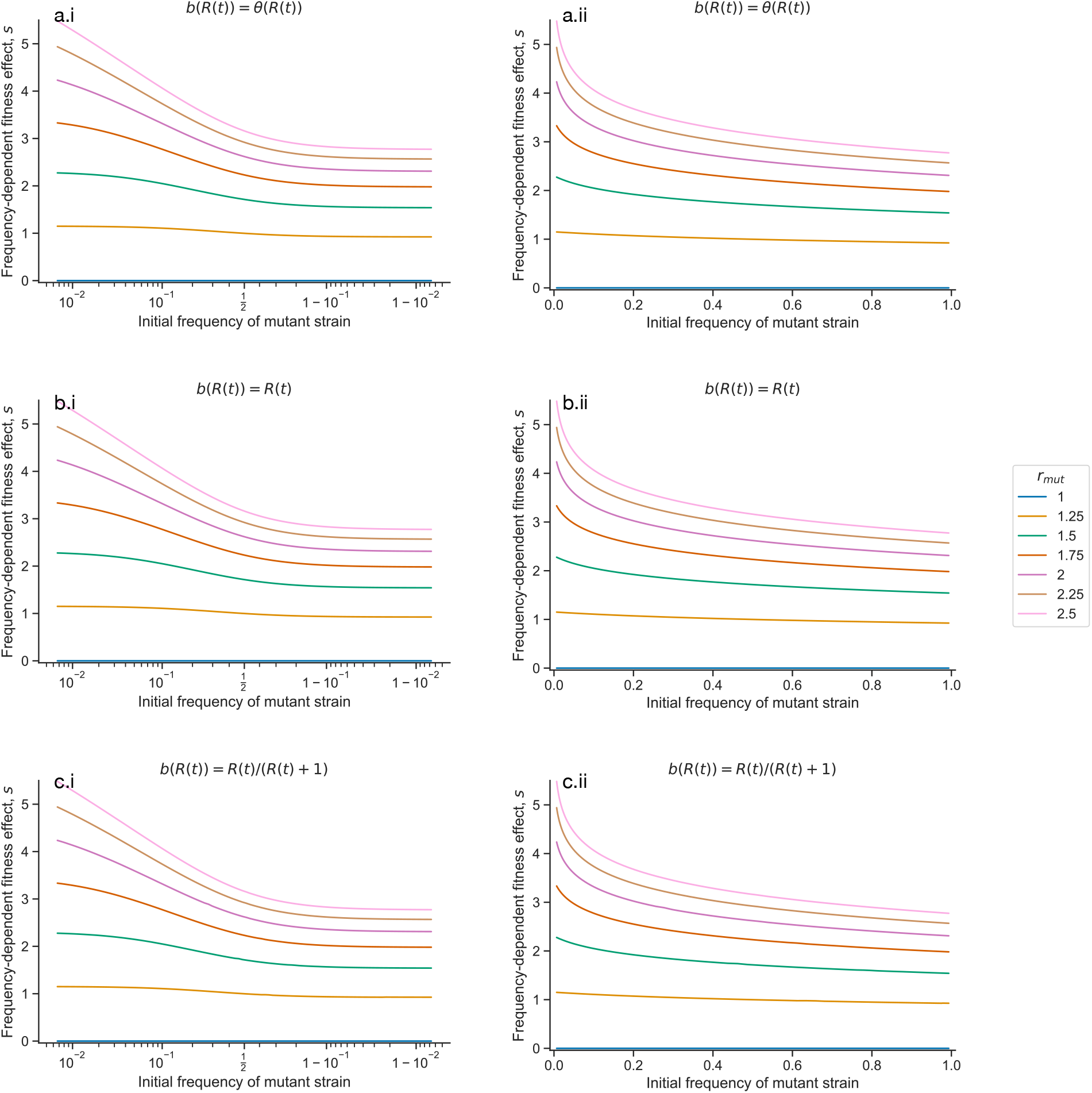
Frequency-dependence arising from resource competition dynamics. Here we show examples of the effective negative frequency-dependent fitness effects that arise from batch culture resource competition dynamics, obtained by numerically solving equations 2-4 (in the main text). We choose three different values of *b*(*R*(*t*)), (a) a step function, (b) a linear function, and (c) a hill function. We show the same plots on either a (i) logit-scaled x-axis, or a (ii) linear-scaled x-axis. Parameter values: wild-type growth rate, *r*_*wt*_ = 1, initial amount of resource, *R*_0_ = 1; initial total population size *N* = 10^−2^; *a*_*wt*_ = *a*_*mut*_ = 1.

**Figure S19:**
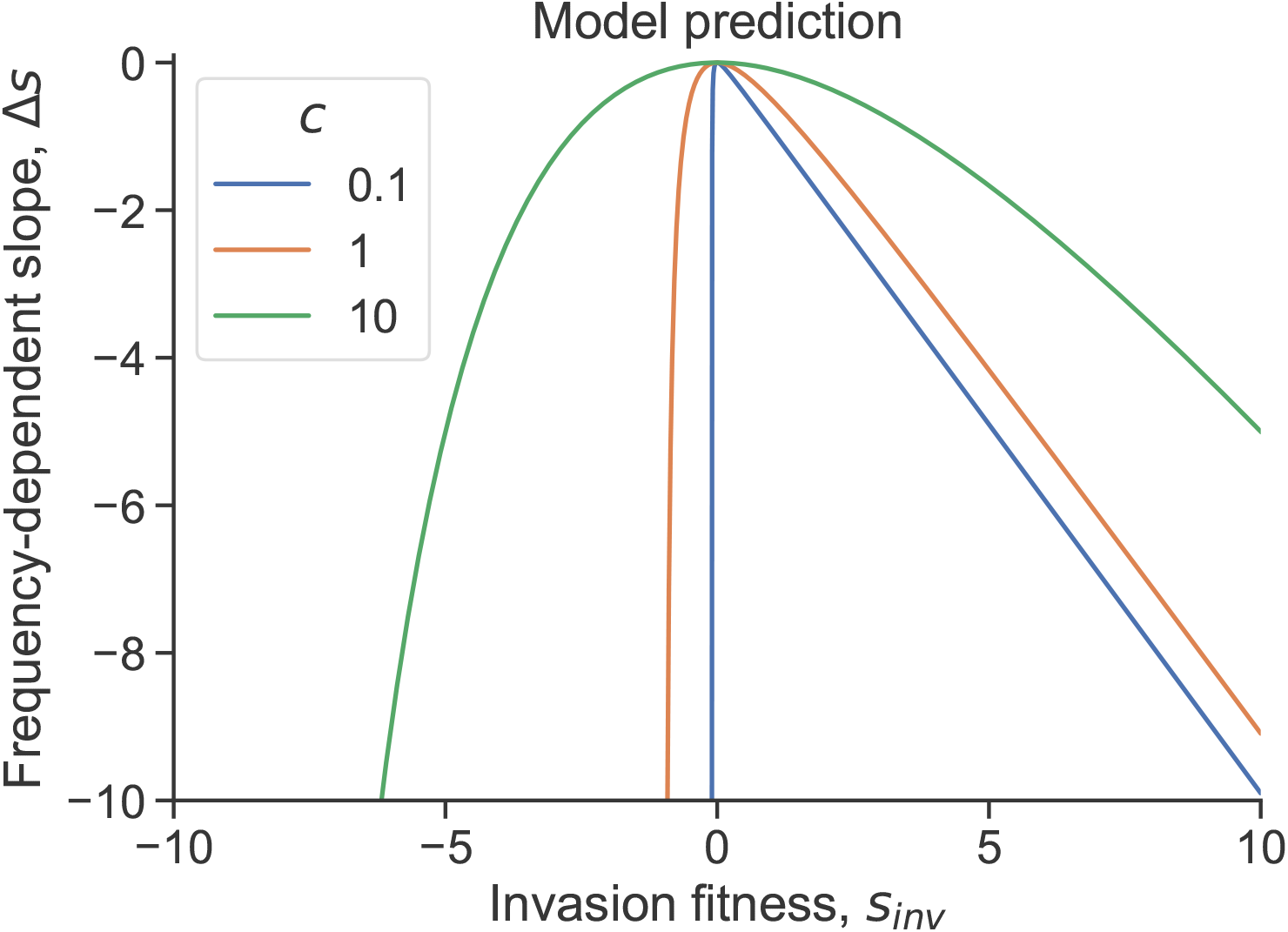
Frequency-dependent slopes predicted from resource competition dynamics. Prediction for the frequency-dependent slope from equation S54, varying *c*. We held *r*_*wt*_ = 1 constant and *a*_*wt*_ = *a*_*mut*_ = 1, and varied *r*_*mut*_ to change *s*_*inv*_.

**Figure S20:**
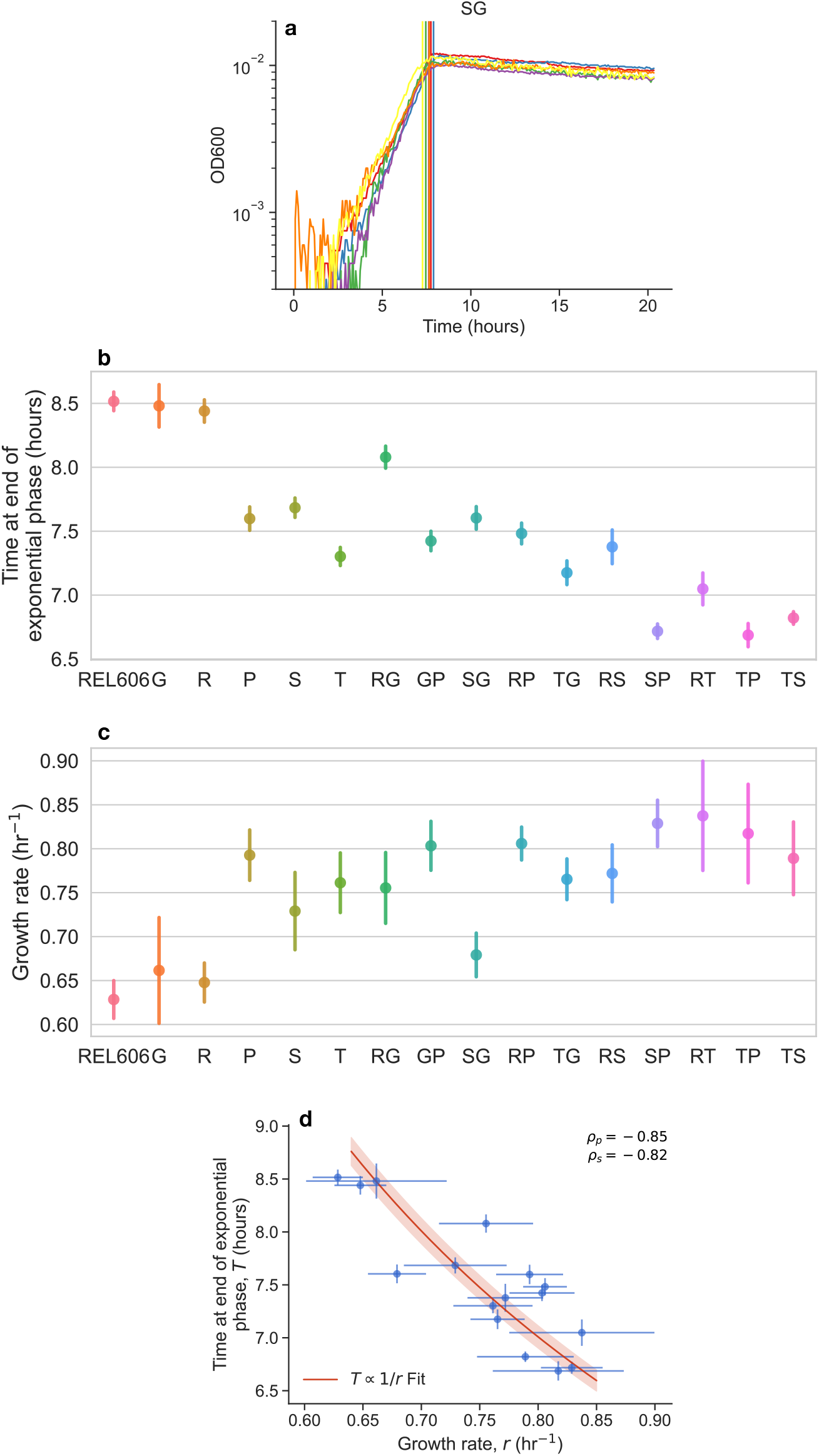
Growth curves of monocultures from plate reader. We measured growth curves using a plate reader of all 16 strains (six biological replicates per strain); (a) example of growth curves, along with estimated time at the end of exponential phase. We estimated (b) the time at the end of exponential phase (*T* ) and (c) average growth rate (*r*) for each strain. (d) We see a strong negative correlation between *r* and and *T*, consistent with a *T* ∝ 1*/r* relationship. All error bars represent standard errors. *ρ*_*p*_ represents Pearson’s correlation; *ρ*_*s*_ represents Spearman’s correlation.

## References

1. Lenski, R. E. Experimental evolution and the dynamics of adaptation and genome evolution in microbial populations. The ISME J. 11, 2181–2194, DOI:10.1038/ismej.2017.69 (2017).

2. Haldane, J. B. S. A Mathematical Theory of Natural and Artificial Selection, Part V: Selection and Mutation. Math. Proc. Camb. Philos. Soc. 23, 838–844, DOI:10.1017/S0305004100015644 (1927).

3. Fisher, R. The Genetical Theory of Natural Selection (1930). Cc: us Lang: en Tab: overview.

4. Kimura, M. The Neutral Theory of Molecular Evolution (Cambridge University Press, Cambridge, 1983).

5. Gillespie, J. H. Molecular Evolution Over the Mutational Landscape. Evolution 38, 1116, DOI:10.2307/2408444 (1984).

6. Orr, H. A. The Population Genetics of Adaptation: The Distribution of Factors Fixed during Adaptive Evolution. Evolution 52, 935, DOI:10.2307/2411226 (1998).

7. Eyre-Walker, A. & Keightley, P. D. The distribution of fitness effects of new mutations. Nat. Rev. Genet. 8, 610–618, DOI:10.1038/nrg2146 (2007).

8. Bataillon, T. & Bailey, S. F. Effects of new mutations on fitness: insights from models and data. Annals New York Acad. Sci. 1320, 76–92, DOI:10.1111/nyas.12460 (2014). _eprint: https://nyaspubs.onlinelibrary.wiley.com/doi/pdf/10.1111/nyas.12460.

9. Orr, H. A. The distribution of fitness effects among beneficial mutations. Genetics 163, 1519 (2003).

10. Good, B. H., Martis, S. & Hallatschek, O. Adaptation limits ecological diversification and promotes ecological tinkering during the competition for substitutable resources. Proc. Natl. Acad. Sci. 115, DOI:10.1073/pnas.1807530115 (2018).

11. Kimura, M. The Number of Heterozygous Nucleotide Sites Maintained in a Finite Population Due to Steady Flux of Mutations. Genetics 61, 893–903 (1969).

12. Ewens, W. J. Mathematical Population Genetics 1: Theoretical Introduction (Springer Science & Business Media, 2004).

13. Crow, J. F. & Kimura, M. An Introduction to Population Genetics Theory (Blackburn Press, 2009). Google-Books-ID: VWqKPwAACAAJ.

14. Desai, M. M. & Fisher, D. S. Beneficial Mutation–Selection Balance and the Effect of Linkage on Positive Selection. Genetics 176, 1759–1798, DOI:10.1534/genetics.106.067678 (2007).

15. Elena, S. F. & Lenski, R. E. Evolution experiments with microorganisms: the dynamics and genetic bases of adaptation. Nat. Rev. Genet. 4, 457–469, DOI:10.1038/nrg1088 (2003).

16. Wiser, M. J., Ribeck, N. & Lenski, R. E. Long-term dynamics of adaptation in asexual populations. Sci. (New York, N.Y.) 342, 1364–1367, DOI:10.1126/science.1243357 (2013).

17. Lang, G. I. et al. Pervasive genetic hitchhiking and clonal interference in forty evolving yeast populations. Nature 500, 571–574, DOI:10.1038/nature12344 (2013).

18. Frean, M. & Abraham, E. R. Rock-scissors-paper and the survival of the weakest. Proc. Royal Soc. B: Biol. Sci. 268, 1323–1327, DOI:10.1098/rspb.2001.1670 (2001).

19. Kerr, B., Riley, M. A., Feldman, M. W. & Bohannan. B. J. Local dispersal promotes biodiversity in a real-life game of rock–paper–scissors. Nature 418, 171–174, DOI:10.1038/nature00823 (2002).

20. Bajić, D., Vila, J. C. C., Blount, Z. D. & Sánchez, A. On the deformability of an empirical fitness landscape by microbial evolution. Proc. Natl. Acad. Sci. United States Am. 115, 11286–11291, DOI:10.1073/pnas.1808485115 (2018).

21. Ascensao, J. A., Wetmore, K. M., Good, B. H., Arkin, A. P. & Hallatschek, O. Quantifying the local adaptive landscape of a nascent bacterial community. Nat. Commun. 14, 248, DOI:10.1038/s41467-022-35677-5 (2023). Number: 1.

22. Le Gac, M., Plucain, J., Hindré, T., Lenski, R. E. & Schneider, D. Ecological and evolutionary dynamics of coexisting lineages during a long-term experiment with Escherichia coli. Proc. Natl. Acad. Sci. United States Am. 96, 10242–7, DOI:10.1073/pnas.96.18.10242 (2012).

23. Post, D. M. & Palkovacs, E. P. Eco-evolutionary feedbacks in community and ecosystem ecology: interactions between the ecological theatre and the evolutionary play. Philos. transactions Royal Soc. London. Ser. B, Biol. sciences 364, 1629–40, DOI:10.1098/rstb.2009.0012 (2009).

24. Laland, K. N., Odling-Smee, F. J. & Feldman, M. W. Evolutionary consequences of niche construction and their implications for ecology. Proc. Natl. Acad. Sci. 96, 10242–10247, DOI:10.1073/pnas.96.18.10242 (1999).

25. Harrington, K. I. & Sanchez, A. Eco-evolutionary dynamics of complex social strategies in microbial communities. Commun. & Integr. Biol. 7, e28230, DOI:10.4161/cib.28230 (2014).

26. Venkataram, S., Kuo, H.-Y., Hom, E. F. Y. & Kryazhimskiy, S. Mutualism-enhancing mutations dominate early adaptation in a microbial community. bioRxiv 2021.07.07.451547, DOI:10.1101/2021.07.07.451547 (2022).

27. Ascensao, J. A. et al. Rediversification following ecotype isolation reveals hidden adaptive potential. Curr. Biol. 34, 855–867.e6, DOI:10.1016/j.cub.2024.01.029 (2024).

28. Yan, L., Neher, R. A. & Shraiman, B. I. Phylodynamic theory of persistence, extinction and speciation of rapidly adapting pathogens. eLife 8, DOI:10.7554/elife.44205 (2019).

29. King, K. C., Delph, L. F., Jokela, J. & Lively, C. M. The geographic mosaic of sex and the Red Queen. Curr. biology: CB 19, 1438–1441, DOI:10.1016/j.cub.2009.06.062 (2009).

30. Morran, L. T., Schmidt, O. G., Gelarden, I. A., Parrish, R. C. & Lively, C. M. Running with the Red Queen: host-parasite coevolution selects for biparental sex. Sci. (New York, N.Y.) 333, 216–218, DOI:10.1126/science.1206360 (2011).

31. Lenski, R. E. Revisiting the Design of the Long-Term Evolution Experiment with Escherichia coli. J. Mol. Evol. 91, 241–253, DOI:10.1007/s00239-023-10095-3 (2023).

32. Brockhurst, M. A., Colegrave, N., Hodgson, D. J. & Buckling, A. Niche occupation limits adaptive radiation in experimental microcosms. PLoS One 2, e193 (2007).

33. Ascensao, J. A. & Desai, M. M. Experimental evolution in an era of molecular manipulation. Nat. Rev. Genet. 1–15, DOI:10.1038/s41576-025-00867-6 (2025).

34. RB, H., CN, V. & J, A. Evolution of Escherichia coli during growth in a constant environment. Genetics 116, 349–358, DOI:10.1093/GENETICS/116.3.349 (1987).

35. Rosenzweig, R. F., Sharp, R. R., Treves, D. S. & Adams, J. Microbial evolution in a simple unstructured environment: genetic differentiation in Escherichia coli. Genetics 137, 903–917, DOI:10.1093/genetics/137.4.903 (1994).

36. Treves, D. S., Manning, S. & Adams, J. Repeated evolution of an acetate-crossfeeding polymorphism in long-term populations of Escherichia coli. Mol. Biol. Evol. 15, 789–797, DOI:10.1093/oxfordjournals.molbev.a025984 (1998).

37. Turner, P. E., Souza, V. & Lenski, R. E. Tests of ecological mechanisms promoting the stable coexistence of two bacterial genotypes. Ecology 77, 2119–2129, DOI:10.2307/2265706 (1996).

38. Rainey, P. B. & Travisano, M. Adaptive radiation in a heterogeneous environment. Nature 394, 69–72, DOI: 10.1038/27900 (1998).

39. Frenkel, E. M. et al. Crowded growth leads to the spontaneous evolution of semistable coexistence in laboratory yeast populations. Proc. Natl. Acad. Sci. 112, 11306–11311, DOI:10.1073/pnas.1506184112 (2015).

40. Elena, S. F. & Lenski, R. E. LONG-TERM EXPERIMENTAL EVOLUTION IN ESCHERICHIA COLI. VII. MECHANISMS MAINTAINING GENETIC VARIABILITY WITHIN POPULATIONS. Evol. Int. J. Org. Evol. 51, 1058–1067, DOI:10.1111/j.1558-5646.1997.tb03953.x (1997).

41. Rozen, D. E. & Lenski, R. E. Long-Term Experimental Evolution in Escherichia coli. VIII. Dynamics of a Balanced Polymorphism. The Am. naturalist 155, 24–35, DOI:10.1086/303299 (2000).

42. Plucain, J. et al. Epistasis and allele specificity in the emergence of a stable polymorphism in Escherichia coli. Science 343, 1366–1369, DOI:10.1126/science.1248688 (2014).

43. Good, B. H., McDonald, M. J., Barrick, J. E., Lenski, R. E. & Desai, M. M. The dynamics of molecular evolution over 60,000 generations. Nature 551, 45–50, DOI:10.1038/nature24287 (2017).

44. Maddamsetti, R., Lenski, R. E. & Barrick, J. E. Adaptation, clonal interference, and frequency-dependent interactions in a long-term evolution experiment with Escherichia coli. Genetics 200, 619–631, DOI:10.1534/genetics.115.176677 (2015).

45. Peng, F. et al. Effects of Beneficial Mutations in pykF Gene Vary over Time and across Replicate Populations in a Long-Term Experiment with Bacteria. Mol. Biol. Evol. 35, 202–210, DOI:10.1093/molbev/msx279 (2018).

46. Khan, A. I., Dinh, D. M., Schneider, D., Lenski, R. E. & Cooper, T. F. Negative Epistasis Between Beneficial Mutations in an Evolving Bacterial Population. Science 332, 1193–1196, DOI:10.1126/science.1203801 (2011).

47. Zhao, S. et al. Adaptive Evolution within Gut Microbiomes of Healthy People. Cell Host Microbe 25, 656–667, DOI:10.1016/j.chom.2019.03.007 (2019).

48. Folkesson, A. et al. Adaptation of Pseudomonas aeruginosa to the cystic fibrosis airway: An evolutionary perspective, DOI:10.1038/nrmicro2907 (2012).

49. Sousa, A. et al. Recurrent Reverse Evolution Maintains Polymorphism after Strong Bottlenecks in Commensal Gut Bacteria. Mol. Biol. Evol. 34, 2879–2892, DOI:10.1093/MOLBEV/MSX221 (2017).

50. Garud, N. R., Good, B. H., Hallatschek, O. & Pollard, K. S. Evolutionary dynamics of bacteria in the gut microbiome within and across hosts. bioRxiv 1–38 (2017).

51. Roodgar, M. et al. Longitudinal linked-read sequencing reveals ecological and evolutionary responses of a human gut microbiome during antibiotic treatment. Genome Res. 31, 1433–1446, DOI:10.1101/gr.265058.120 (2021).

52. Schlechter, R. O. et al. Chromatic Bacteria - A Broad Host-Range Plasmid and Chromosomal Insertion Toolbox for Fluorescent Protein Expression in Bacteria. Front. microbiology 9, DOI:10.3389/FMICB.2018.03052 (2018).

53. Kinsler, G., Geiler-Samerotte, K. & Petrov, D. A. Fitness variation across subtle environmental perturbations reveals local modularity and global pleiotropy of adaptation. eLife 9, e61271, DOI:10.7554/eLife.61271 (2020).

54. Kryazhimskiy, S., Rice, D. P., Jerison, E. R. & Desai, M. M. Global epistasis makes adaptation predictable despite sequence-level stochasticity. Sci. (New York, N.Y.) 344, 1519–1522, DOI:10.1126/science.1250939 (2014).

55. Manhart, M., Adkar, B. V. & Shakhnovich, E. I. Trade-offs between microbial growth phases lead to frequency-dependent and non-transitive selection. Proc. Royal Soc. B: Biol. Sci. 285, 20172459, DOI:10.1098/rspb.2017.2459 (2018).

56. Lin, J., Manhart, M. & Amir, A. Evolution of Microbial Growth Traits Under Serial Dilution. Genetics 215, 767–777, DOI:10.1534/genetics.120.303149 (2020).

57. Feng, Z., Blumenthal, E., Mehta, P. & Goyal, A. A theory of ecological invasions and its implications for eco-evolutionary dynamics. Proc. Natl. Acad. Sci. 122, e2505850122, DOI:10.1073/pnas.2505850122 (2025).

58. Li, S. Y., Feng, Z., Goyal, A. & Mehta, P. Emergent frequency-dependent selection predicts mutation outcomes in complex ecological communities, DOI:10.48550/arXiv.2509.23977 (2026). ArXiv:2509.23977 [q-bio.PE].

59. Kinsler, G., Li, Y., Sherlock, G. & Petrov, D. A. A high-resolution two-step evolution experiment in yeast reveals a shift from pleiotropic to modular adaptation. PLOS Biol. 22, e3002848, DOI:10.1371/journal.pbio.3002848 (2024).

60. Mukherjee, A. et al. Coexisting ecotypes in long-term evolution emerged from interacting trade-offs. Nat. Commun. 14, 3805, DOI:10.1038/s41467-023-39471-9 (2023). Number: 1.

61. Scott, M., Gunderson, C. W., Mateescu, E. M., Zhang, Z. & Hwa, T. Interdependence of Cell Growth and Gene Expression: Origins and Consequences. Science 330, 1099–1102, DOI:10.1126/science.1192588 (2010).

62. Basan, M. et al. Overflow metabolism in Escherichia coli results from efficient proteome allocation. Nature 528, 99–104, DOI:10.1038/nature15765 (2015).

63. Basan, M. et al. A universal trade-off between growth and lag in fluctuating environments. Nature 584, 470–474, DOI:10.1038/s41586-020-2505-4 (2020).

64. Vasi, F., Travisano, M. & Lenski, R. E. Long-Term Experimental Evolution in Escherichia coli. II. Changes in Life-History Traits During Adaptation to a Seasonal Environment. https://doi.org/10.1086/285685 144, 432–456, DOI:10.1086/285685 (1994).

65. Orr, H. A. The genetic theory of adaptation: a brief history. Nat. Rev. Genet. 6, 119–127, DOI:10.1038/nrg1523 (2005).

66. de Visser, J. A. G. M., Cooper, T. F. & Elena, S. F. The causes of epistasis. Proc. Royal Soc. B: Biol. Sci. 278, 3617–3624, DOI:10.1098/rspb.2011.1537 (2011).

67. Pavlicev, M. & Wagner, G. P. A model of developmental evolution: selection, pleiotropy and compensation. Trends Ecol. & Evol. 27, 316–322, DOI:10.1016/j.tree.2012.01.016 (2012).

## References

[1] Joao A. Ascensao, Jonas Denk, Kristen Lok, QinQin Yu, Kelly M. Wetmore, and Oskar Hallatschek. Re-diversification following ecotype isolation reveals hidden adaptive potential. Current Biology, 34(4):855–867.e6, February 2024.

[2] Rudolf O. Schlechter, Hyunwoo Jun, Michał Bernach, Simisola Oso, Erica Boyd, Dian A. Muñoz-Lintz, Renwick C.J. Dobson, Daniela M. Remus, and Mitja N.P. Remus-Emsermann. Chromatic Bacteria-A Broad Host-Range Plasmid and Chromosomal Insertion Toolbox for Fluorescent Protein Expression in Bacteria. Frontiers in microbiology, 9(DEC), 12 2018.

